# The aphid effector CathB6 suppresses EDS1-dependent immunity and rewires MORF2–GUN1–GLK signalling to sustain host cellular homeostasis in Arabidopsis

**DOI:** 10.64898/2026.08.10.743731

**Authors:** Qun Liu, Sam T. Mugford, Jianhua Huang, Anna C. M. Neefjes, Saskia A. Hogenhout

## Abstract

Aphids use stylet mouthparts to probe plant tissues and deliver oral secretions into host cells, modulating immunity to establish long-term phloem feeding sites in sieve elements. This feeding behaviour make aphids efficient vectors of plant pathogens. We previously showed that *Myzus persicae* cathepsin B (CathB) effectors promote aphid colonization, accumulate in dynamic cytoplasmic processing bodies (p-bodies), bind the key *Arabidopsis thaliana* immune regulator EDS1, and recruit EDS1 together with its signalling partners PAD4 and ADR1 to p-bodies. However, the extent to which CathB effectors suppress EDS1-regulated immunity remains unclear. Here, we show that CathB6-expressing Arabidopsis lines exhibit substantial transcriptomic overlap with the *eds1-2* mutant, consistent with CathB6 suppression of EDS1-dependent immunity. However, CathB6 also induces broader transcriptional reprogramming beyond that explained by loss of EDS1 function. A yeast two-hybrid screen identified the transcription factors GLK1 and GLK2, as well as several MORF proteins, as CathB6 interactors. CathB6-expressing plants recapitulated GLK- and MORF-regulated transcriptional changes, suppressing GUN1-associated activity while maintaining GLK-mediated expression of photosynthesis-associated nuclear genes (PhANGs). We found that GLKs promotes CathB6 nuclear accumulation and reduces its p-body association. TurboID proximity labelling further linked CathB6 to PhANG-associated proteins, known MORF2- and GUN1-interacting factors, p-body components, and actin–tubulin/myosin machinery, the latter being consistent with the highly dynamic behaviour of CathB6-associated p-bodies. Together, these data indicate that, beyond suppressing EDS1-mediated defenses, CathB6 interferes with the MORF2–GUN1–GLK signalling pathway, modulating plant defense responses while sustaining GLK-mediated PhANG expression to maintain cellular homeostasis during aphid attack.

**SIGNIFICANCE STATEMENT:** Aphids damage crops and spread plant diseases while feeding. We discovered how a protein in aphid saliva, CathB6, helps these pests establish themselves on plants. CathB6 weakens a major immune pathway and alters communication between chloroplasts and the cell nucleus. This allows the aphid to reduce plant defences while maintaining photosynthesis and normal cell function. These findings reveal a sophisticated survival strategy and may identify ways to develop crops that are more resistant to aphids.

## INTRODUCTION

Aphids are highly specialized, plant-dependent insects that have co-evolved with their hosts for approximately 250 million years. These phloem feeders use stylet mouthparts to navigate between plant cells, repeatedly probing individual cells and injecting oral secretions into the cytoplasm before sampling cellular contents (Tjallingii and Esch, 1993). This pre-phloem probing continues until the stylets reach sieve elements, where aphids establish long-term feeding sites. Such intimate cellular interactions make aphids powerful manipulators of plant physiology and highly efficient vectors of plant pathogens, particularly viruses (Martín et al., 1997).

Why aphids engage in extensive pre-phloem probing remains unclear. One possibility is that salivary secretions delivered during this process progressively reprogramme host immune signalling to facilitate stable phloem feeding (Giolai, 2019). However, major gaps remain in defining the effector proteins of the green peach aphid *Myzus persicae* (Sulzer), their host targets, and the molecular pathways through which they reshape host immune and cellular states (Nalam et al., 2013; Pitino and Hogenhout, 2013; Gravino et al., 2025; Liu et al., 2025).

*M. persicae* is particularly well suited for dissecting these mechanisms due to its exceptional versatility as a generalist, colonizing over 400 plant species across monocots and dicots, despite its predominantly clonal reproduction (CABI, 2022). Individuals within one clonal lineage show pronounced host-dependent transcriptional plasticity (Mathers et al., 2017; Chen et al., 2020). This plasticity includes differential expression of genes within an expanded clade of the cathepsin B (CathB) family, including being upregulated in aphids that colonize *Arabidopsis thaliana* and *Brassica rapa* compared to other plant species, implicating these proteins in host adjustment (Mathers et al., 2017; Chen et al., 2020; Liu et al., 2025). Consistent with this, multiple peptides corresponding to the CathB proteins within this expanded clade were detected in aphid oral secretions, unlike other CathB proteins. Moreover, *A. thaliana* lines that stably produce individual CathB proteins within the expanded clade are more susceptibe to *M. persicae* (Liu et al., 2025).

The CathB protein family has followed an unusual evolutionary trajectory in aphids. Similarly to CathB proteins in *M. persicae*, earlier reports provided evidence that CathB family members have expanded and undergone positive selection pressures in the pea aphid *Acyrthosiphon pisum* as well as in several social aphid species of the *Tuberaphis* genus (Kutsukake et al., 2004; Kutsukake et al., 2008; Rispe et al., 2008). Interestingly, soldier aphids of the latter have co-opted CathB as a venom, which is injected into predators via their stylets inducing rapid paralysis (Kutsukake et al., 2004; McKerrow et al., 2006). CathB proteases have important roles in lysosome functions of animal cells and protein turnover and cellular homeostasis in plant cells (Foghsgaard et al., 2001; Carter et al., 2004; Cavallo-Medved et al., 2011; Aggarwal and Sloane, 2014; Bárány et al., 2018). However, protease activity is not required for *M. persicae* CathB proteins within the expanded clade to promote plant susceptibility to aphids, suggesting functional diversification, and leading to the conclusion that CathB proteins of plant-feeding aphids are moonlighting proteases that have evolved as effectors, in agreement with recent findings that *M. persicae* CathB proteins recruit proteins with key roles in regulating plant defence to processing bodies (p-bodies) (Liu et al., 2025).

P-bodies are membrane-less RNA–protein condensates involved in mRNA storage, translational repression and RNA turnover (Brengues et al., 2005; Jang et al., 2019; Kearly et al., 2024; Liu et al., 2025). *M. persicae* CathB effectors directly interact with *A. thaliana* ENHANCED DISEASE SUSCEPTIBILITY 1 (EDS1, AT3G48090), a central immune regulator, and recruit EDS1 together with its signalling partners PHYTOALEXIN DEFICIENT 4 (PAD4, AT3G52430) and ACTIVATED DISEASE RESISTANCE 1 (ADR1, AT1G33560) to p-bodies (Liu et al., 2025). This plant immune complex promotes resistance to aphids, though aphid fecundity increases less markedly on *A. thaliana eds1* mutants than on *pad4* or *adr1* triple mutants (Pegadaraju et al., 2007; Dongus et al., 2022; Liu et al., 2025), while expression of the PAD4 lipase domain alone is sufficient to enhance aphid resistance (Dongus et al., 2019). These observations may reflect the highly effective suppression of EDS1 activity by aphids (Liu et al., 2025). CathB-mediated sequestration of EDS1 complexes to p-bodies could therefore attenuate plant defense to aphids defense, consistent with emerging evidence linking p-bodies to immunity (Bach-Pages et al., 2025). This model is supported by the observation that the *A. thaliana* Hsp20 protein Acd28.9 (AT5G47590) binds CathB effectors, antagonizes their p-body accumulation and EDS1 complex recruitment, and restricts aphid susceptibility (Liu et al., 2025). Conversely, a CathB propeptide activates defense in *Nicotiana tabacum* partly by downregulating EDR1- like suppressors (Guo et al., 2020). Together, these findings suggest that expanded aphid CathB clade act as salivary effectors modulating plant immunity in a host-specific manner, but the downstream pathways they target remain largely unknown.

Here, we investigated the extent to which the *M. persicae* effector CathB6 alters responses in *A. thaliana*. Transcriptomic comparisons revealed that CathB6 expression produces a signature partially resembling the *eds1-2* immune mutant, consistent with the suppression of EDS1-dependent immunity. However, CathB6 induced broader transcriptional changes beyond those explained by EDS1 loss alone, indicating additional host targets. Interaction and proximity-labelling datasets linked CathB6 to MULTIPLE ORGANELLAR RNA EDITING FACTOR (MORF) proteins, GOLDEN2-LIKE (GLK) transcription factors, components of GENOMES UNCOUPLED 1 (GUN1)-mediated chloroplast-to-nucleus signalling, and proteins involved in organellar RNA editing, translation and cytoplasmic trafficking. Together, our data suggest that CathB6 perturbs the MORF–GUN1–GLK pathway to suppress SA-related immunity while sustaining GLK-driven expression of photosynthesis-associated nuclear genes (PhANGs), revealing a molecular mechanism by which an aphid salivary effector modulates both host immunity and cellular homeostasis during colonization.

## RESULTS

### CathB6 effects on the plant transcriptome overlap with EDS1-regulated immunity

To asses if CathB6 suppresses plant immune responses regulated by EDS1, we conducted RNA-seq analyses of CathB6-GFP transgenic and *eds1-2* mutant lines. Principal component analysis (PCA) with the RNA-seq data revealed that wild-type and GFP-expressing control plants separated from both CathB6-GFP and *eds1-2* samples along PC1, which explained 46% of the variance, indicating that the major source of transcriptional variance is shared between CathB6 expression and EDS1 deficiency (Figure 1A). Analysis of differentially expressed (DE) transcripts confirmed this relationship (Table S1). A Venn diagram showed a distinct and statistically significant overlap of DE transcripts between the CathB6-GFP and *eds1-2* datasets (*P* < 0.01, Fisher’s exact test), with 42 genes (3.4%) upregulated in both CathB6-GFP transgenic and *eds1-2* mutant lines and 92 genes (7.5%) downregulated in both lines (Figure 1B). Hierarchical clustering demonstrated substantial co-regulation of the shared transcriptome (Figure 1C).

**Figure 1.**
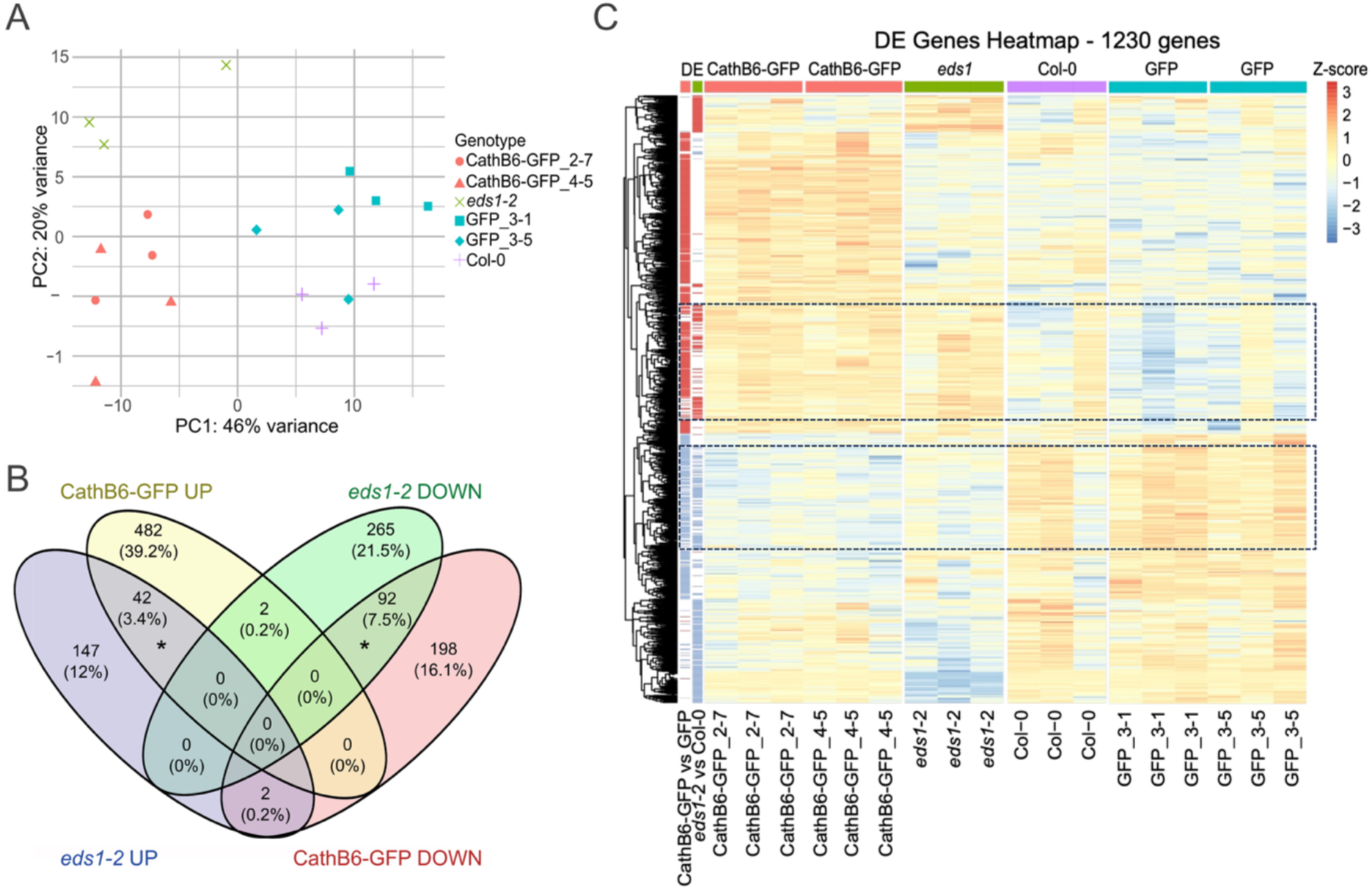
The transcriptional profiles of *A. thaliana* CathB6-GFP and *eds1-2* plants overlap. (A) Principal component analysis of RNA-seq read counts. PC1 separates CathB6-GFP and *eds1-2* samples from GFP-expressing and wild-type Col-0 plants, indicating shared variance. PC2 further distinguishes CathB6-GFP from *eds1-2*, revealing divergence between the two genotypes. (B) Venn diagram showing the overlap of differentially expressed (DE) transcripts between CathB6-GFP (vs. GFP) and eds1-2 (vs. Col-0). The asterisks indicate statistically significant overlaps (P < 0.01, Fisher’s exact test). (C) Hierarchical clustering (dendrogram on the left) of DE transcripts (heatmap on the right, red = up-regulated and blue = down-regulated). The clustering separates transcripts with similar regulation by CathB6-GFP and EDS1 from those exhibiting CathB6-GFP-specific or EDS1-specific regulation. Dashed boxes indicate clusters of transcripts that are co-regulated in *A. thaliana* CathB6-GFP and *eds1-2* plants.

Gene ontology (GO) term enrichment analysis (Fig.S1; Tables S2 and S3) revealed 33 GO categories enriched for dowregulated genes in the *eds1-2* vs Col-0 wild-type lines, including those involved in plant responses to various stimuli, plant defence and cell death, and responses to oxidative stress, of which the GO category response to singlet oxygen (GO:0000304) was the most enriched, in line with the role of EDS1 in regulating plant immunity (Wiermer et al., 2005). GO terms enriched for upregulated genes in the *eds1-2* vs Col-0 lines totalled 15 and these included responses to hypoxia, oxygen levels, red and far red light and regulation of RNA biosynthetic processes and DNA-templated transcription. The 11 GO terms enriched for downregulated genes in the CathB6-GFP vs GFP lines included responses to hypoxia, oxygen levels, stress and stimulus, and the 8 upregulated GO categories reponses to SA, defence and other organisms. Among proteins that regulate similar processes, upregulated genes between the CathB6-GFP and *eds1-2* lines included those encoding the galactinol synthases GolS2 and GolS4 that promote increased tolerance to salt, chilling and oxidatuve stress (Sun et al., 2013; Song et al., 2016) and downregulated genes those encoding AZELAIC ACID INDUCED 1 (AZI1, AT4G12470), AZI3 (AT4G12490) and EARLI1 (AT4G12480), which are proteins that accumulate at plastids during defence priming and are involved in the priming of salicylic acid induction and systemic immunity triggered by pathogen or azelaic acid (Cecchini et al., 2015) (Table S1). These data are in agreement with the idea that CathB6 modulates processes regulated by EDS1.

Despite substantial overlap, PC2 of the RNA-seq data PCA, explaining 20% of the variance, clearly separated CathB6-GFP from *eds1-2* samples (Figure 1A), indicating transcriptional divergence between the two genotypes. This suggests that a significant portion of CathB6-regulated genes are not mediated by EDS1, pointing to additional, EDS1-independent effects of CathB6 on the plant transcriptome.

### Yeast two-hybrid library screening identifies GLK1, GLK2, MORF2 and Acd28.9 as CathB6 interactors

We investigated what CathB6 may regulate beyond EDS1 in plants, we performed a yeast two-hybrid (Y2H) screen of CathB6 against a cDNA library generated from *A. thaliana* plants exposed to *M. persicae*, other insects and a phytoplasma. This screen yielded 17 candidate interactors from 39 clones (Table S4). We prioritized candidates that appeared in multiple independent clones for further validation. Subsequent independent Y2H assays in two yeast strains, NMY51 (Figure S2A, B) and AH109 (Figure S2C, D), utilizing the LexA and Gal4 DNA-binding domains respectively, confirmed interactions between CathB6 and *A. thaliana* GOLDEN2-LIKE 1 (GLK1, AT2G20570), MULTIPLE ORGANELLAR RNA EDITING FACTOR 2 (MORF2, AT2G33430) and the ALPHA CRYSTALLIN HEAT SHOCK PROTEIN 20 (Hsp20) family protein Acd28.9 (AT5G47590). The latter protein has previously been shown to directly interact with CathB6 and counteract its activity in plants (Liu et al., 2025).

GLK1 and its homolog GLK2 (AT5G44190) are GARP superfamily transcription factors that positively regulate chloroplast development (Fitter et al., 2002). They function redundantly to promote PhANG expression in light, while their expression is repressed by PHYTOCHROME-INTERACTING FACTOR (PIF) transcription factors in darkness (Waters et al., 2009; Martín et al., 2016). Y2H assays indicated that *M. persicae* CathB6 and CathB3 interact with both GLK1 and GLK2 in Y2H experiments, unlike CathB9 (Figure 2A; Figure S2C; Figure S3A), possibly aligning with previous findings that CathB3 and CathB6 are host-responsive and improve aphid fecundity, in contrast to CathB9 (Liu et al., 2025). CathB6-GFP pulled down HA-GLK1 and HA-GLK2, with HA-GLK2 being more abundant than HA-GLK1, in pull-down assays from agroinfiltrated *N. benthamiana* leaves (Figure 2B).

**Figure 2.**
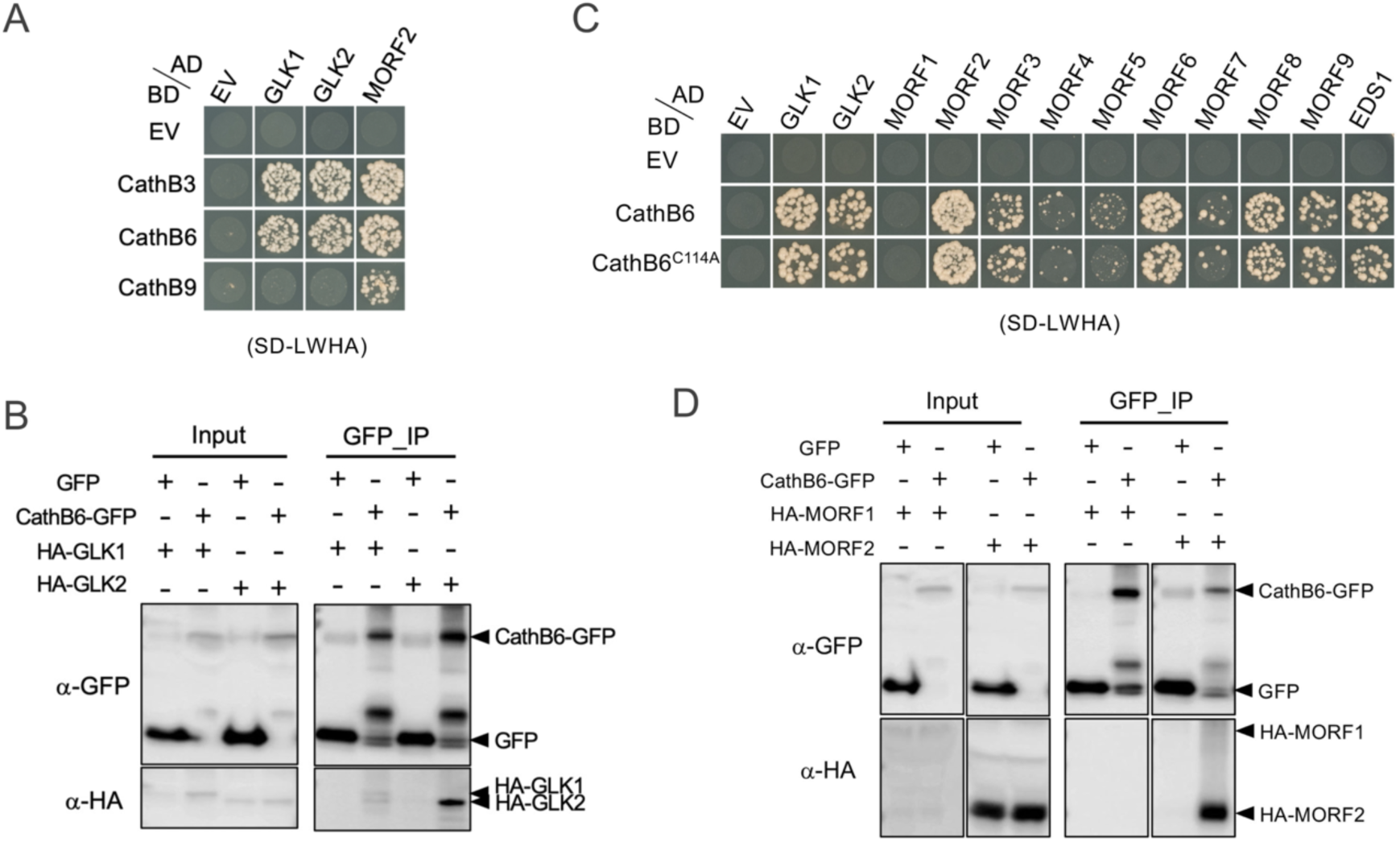
*M. persicae* CathB6 effectors interact with *A. thaliana* GLK and MORF proteins. (A, C) Yeast two-hybrid assays testing interactions between *M. persicae* CathB effectors and members of *A. thaliana* GLK and MORF protein families. CathB6^C114A^ has a mutation in the protease catalytic site. AD, GAL4-activation domain. BD, GAL4-DNA binding domain. SD-LWHA, quadruple dropout medium lacking leucine, tryptophan, histidine, and adenine. Yeast growth on SD-LW medium is shown in Supplementary Fig. 3A, B. (B, D) Co-immunoprecipitation (Co-IP) of HA-tagged GLK and MORF proteins with GFP or CathB6-GFP from agroinfiltrated *N. benthamiana* leaves using GFP-trap beads. Arrowheads indicate the expected sizes: GFP, 26.9 kDa; CathB6-GFP, 59.9 kDa; HA-GLK1, 53.2 kDa; HA-GLK2, 47.4 kDa; HA-MORF1, 50.4 kDa; HA-MORF2, 30.0 kDa.

MORF proteins are required for RNA editing in plastids by forming complexes with pentatricopeptide repeat (PPR) proteins that carry a DWY motif for C-to-U conversion (Takenaka et al., 2012; Yan et al., 2018). Beyond RNA editing, MORF2 also functions in chloroplast-to-nucleus signalling, stress responses and skotomorphogenesis (Yapa et al., 2023; Li et al., 2025). Y2H analyses provided evidence that all three *M. persicae* CathB proteins, CathB3, CathB6 and CathB9, interact with MORF2 (Figure 2A). Moreover, CathB6 interacts with MORF3, MORF6, MORF8 and MORF9, interacts weakly with MORF4, MORF5 and MORF7 and does not interact with MORF1 (Figure 2C; Fig. S3B). CathB6-GFP pulled down HA-MORF2 from agroinfiltrated *N. benthamiana* leaves, whereas HA-MORF1 was not pulled down, though it cannot be excluded that HA-MORF1 was not pulled down due to its lower abundance compared to HA-MORF2 (Figure 2D).

The catalytically inactive CathB6^C114A^ mutant also interacts with GLKs and MORFs (Figure 2C), in agreement with this mutant also promoting aphid fecundity (Liu et al., 2025).

These data indicate that the aphid CathB effectors bind *A. thaliana* GLK and MORF proteins. GLKs controls the expression of photosynthesis-associated nuclear genes (PhANGs), including those encoding MORFs. MORF2 and MORF9 are transported to chloroplasts (Takenaka et al., 2012; Yuan et al., 2022) and MORF1, MORF3, MORF5, MORF6 and MORF8 to both mitochondria and chloroplasts (Takenaka et al., 2012; Glass et al., 2015; Law et al., 2015). Therefore, aphid CathB effectors interact with proteins that regulated nucleus-to-chloroplast communications.

### GLKs and MORF2 impact *M. persicae* fecundity on *A. thaliana*

Next, we conducted aphid fecundity assays on *glk1.1 glk2.1* and *morf2-1 A. thaliana* mutants (Col-0 background, homozygous T-DNA insertion lines). Wild-type Col-0 and GFP transgenic plants were included as controls, along with CathB6-GFP transgenic plants, as well as the *eds1-2* and *acd28.9* mutants, which were previously shown to be more susceptible to aphids (Liu et al., 2025). *M. persicae* reproduction was significantly higher on the *acd28.9* and CathB6-GFP lines compared to Col-0 and GFP controls (Figure 3), confirming previous data (Liu et al., 2025). In contrast, the aphids produced significantly fewer progeny on *glk1.1 glk2.1* mutants relative to Col-0 and all other lines, indicating that GLK1 and GLK2 are required for optimal aphid performance. Aphids performed slightly better on *A. thaliana eds1-2* and *morf2-1* mutants (Figure 3) compared to WT, indicating that MORF2 may negatively impact aphid fecundity.

**Figure 3.**
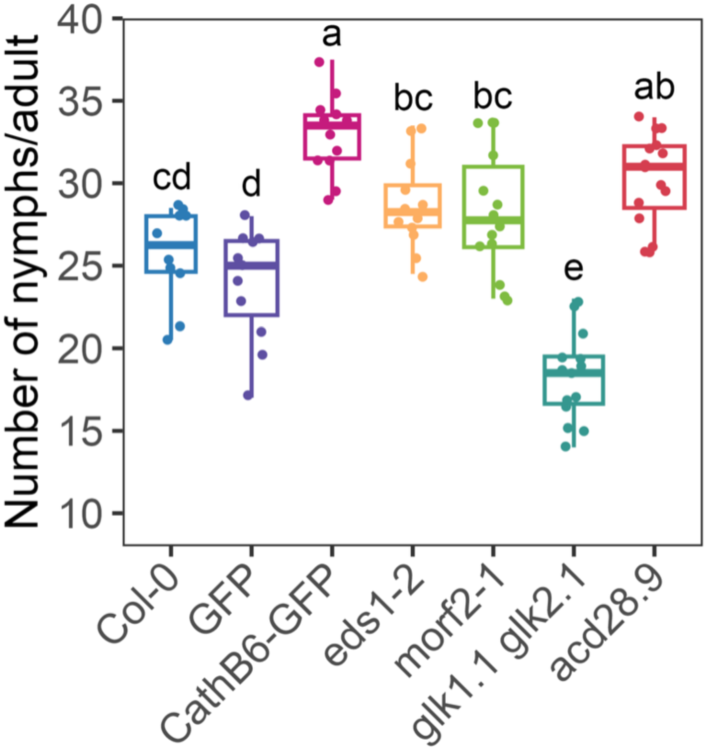
*M. persicae* fecundity assays on *A. thaliana* transgenic and mutant lines. Box plots show fecundity per female aphid at n = 10 to 15 replicates (shown as dots) per line (Col-0, n = 10; GFP, n = 11; CathB6-GFP, n = 12; *eds1-2*, n = 12; *morf2-1*, n = 14; *glk1.1 glk2.1*, n = 14; *acd28.9*, n = 15). Different letters above the boxes indicate significant differences determined by one-way ANOVA with Tukey’s post-hoc test (*P* < 0.05). The transgenic lines GFP (line 3-5) and CathB6-GFP (line 4-5), as well as the mutants *eds1-2* and *acd28.9*, were previously described in Liu et al. (2025). The *morf2-1* (Takenaka et al., 2012) and *glk1.1 glk2.1* (Ahmad et al., 2019) lines were genotyped for this study.

### CathB6 recapitulates GLK- and MORF-regulated transcriptome changes

To examine whether the interactions of CathB6 with MORF2 and GLKs are reflected at the transcriptome level, we compared RNA-seq data from CathB6-GFP plants with DE gene-lists from published datasets from *glk1.1 glk2.1* mutants (Ahmad et al., 2019) and from norflurazon (NF)-treated MORF2-overexpressing (MORF2-OX) and *gun1-9* mutant plants, each compared with NF-treated wild-type controls (Zhao et al., 2019). We included the *gun1-9* mutant in our analyses, because MORF2 interacts with Genomes Uncoupled 1 (GUN1) to regulate plastid RNA editing, while GUN1 contributes to the repression of PhANG expression when plastid function is compromised (Zhao et al., 2019; Wu and Bock, 2021; Loudya et al., 2024). In contrast, GLK transcription factors activate PhANGs required for chloroplast biogenesis (Chen et al., 2016; Li et al., 2022; Zheng et al., 2024). NF inhibits phytoene desaturase (PDS), an early enzyme in carotenoid biosynthesis, causing photooxidative chloroplast damage and leaf or seedling bleaching (Breitenbach et al., 2001). In response to NF-induced plastid dysfunction, GUN1-represses nuclear photosynthesis and chloroplast biogenesis programmes and this repression is impaired in *gun1* mutants, resulting in sustained expression of chloroplast-related nuclear genes despite plastid dysfunction (Zhao et al., 2019). Similarly, MORF2-overexpressing plants show increased expression of nuclear-encoded chloroplast proteins following NF treatment, consistent with disrupted chloroplast-to-nucleus signalling (Zhao et al., 2019).

The comparative analyses if RNAseq data revealed enrichments in gene overlaps between upregulated genes of CathB6-GFP plants and down-regulated genes of *glk1.1 glk2.1* plants (Figure 4A; Table S1). We also identified these enrichments between upregulated genes of CathB6-GFP plants and upregulated genes of NF-treated MORF2-OX plants and downregulated genes between these plants (Figure 4B; Table S1). An even stronger overlap was observed between CathB6-GFP plants and NF-treated *gun1-9* mutants (Figure 4C; Table S1).

**Figure 4.**
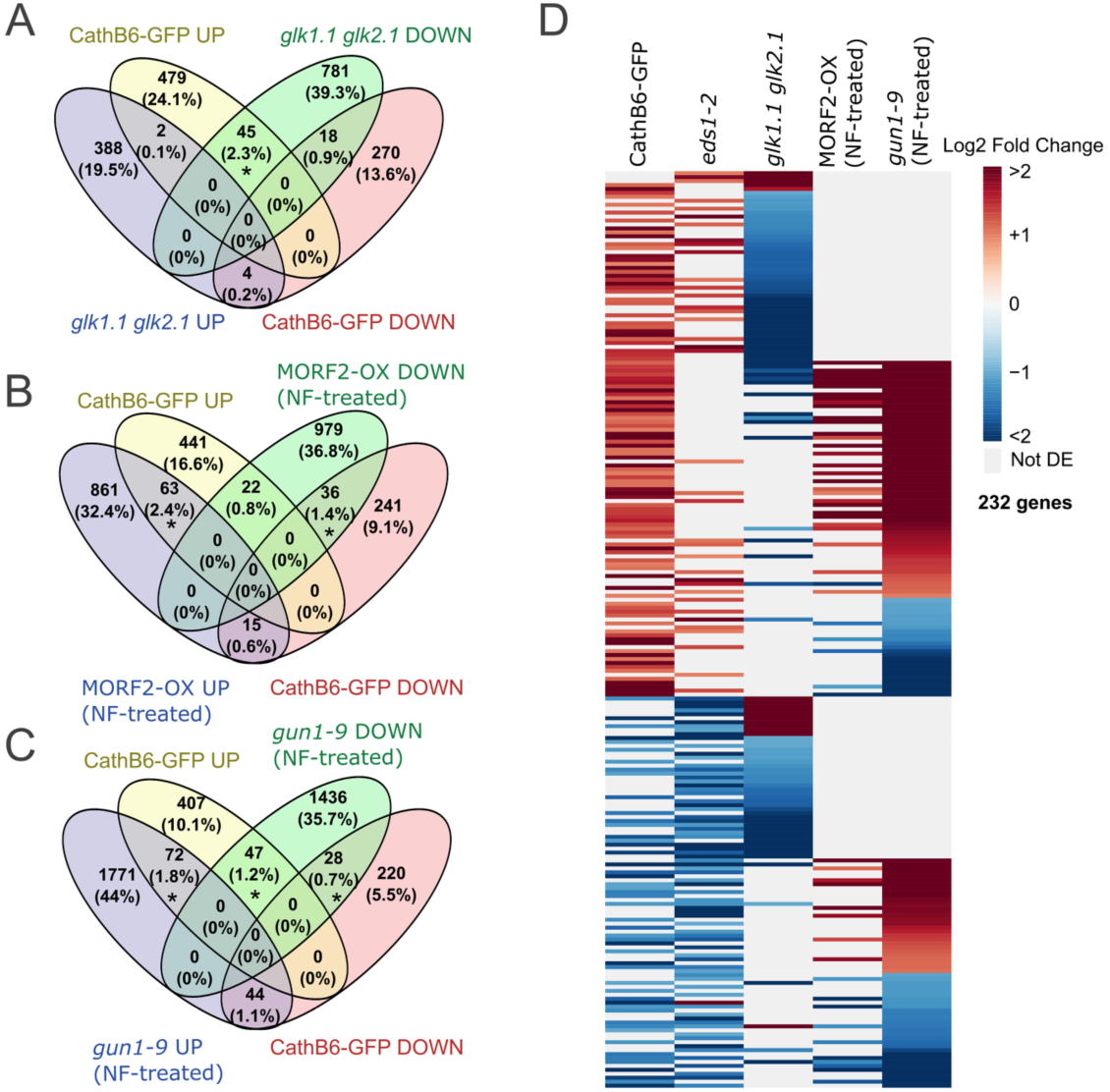
The transcriptional profiles of *A. thaliana* CathB6-GFP plants overlap with those of *glk* and *gun1* mutants and MORF2-ox lines. (A–C) Venn diagrams showing the overlap of genes differentially expressed in CathB6-GFP with those up- (red) or down- (blue) regulated in (A) the *glk1.1 glk2.1* double mutant line (Ahmad et al., 2019) and (B) NF-treated MORF2 overexpression (MORF2-OX) and (C) the NF-treated *gun1-9* mutant lines (Zhao et al., 2019). Asterisks indicate statistically significant overlap with CathB6-GFP-regulated genes (Fisher’s exact test, *P* < 0.05). (D) Heatmap showing relative expression levels of 232 genes that are CathB6-GFP (n= 157) and/or EDS1-regulated (n=112, of which 35 are also CathB6-regulated) and differentially expressed in *glk1.1 glk2.1*, MORF2-OX, or *gun1-9* lines.

Most genes upregulated in CathB6-GFP plants were downregulated in *glk1.1 glk2.1* plants (Figure 4A, D; Table S1). This inverse relationship suggests that CathB6 promotes a subset of GLK-regulated chloroplast genes. In contrast, most genes upregulated in CathB6-GFP plants were also upregulated in NF-treated MORF2-OX and *gun1-9* plants (Figure 4D), though downregulated genes of CathB6-GFP plants showed more variable patterns, with approximately half being upregulated and half downregulated in NF-treated MORF2-OX and *gun1-9* plants (Figure 4D). Most genes differentially regulated in NF-treated MORF2-OX and *gun1-9* plants did not overlap with those altered in *glk1.1 glk2.1* plants. However, the overlapping genes that were identied were predominantly downregulated in *glk1.1 glk2.1* and upregulated in CathB6-GFP, MORF2-OX and *gun1-9* plants (Figure 4D). Together, these data suggest that aphid CathB6 modulates GUN1/MORF2-dependent chloroplast-nucleus signalling in a manner that promotes expression of PhANGs in the nucleus.

### CathB6 reduces editing efficiency of the chloroplast *clpP*-559 transcript

In a well-functioning photosynthetic cell under light conditions, MORF2 regulates efficient RNA editing (Zhao et al., 2019). Conversely, in a dysfunctional photosynthetic cell, GUN1 interacts with MORF2 to suppress editing and maturation of chloroplast-encoded transcripts (Larkin, 2019; Zhao et al., 2019; Tang et al., 2024). However, plastid RNA editing and splicing are not obviously impacted in *gun1* mutants (Tang et al., 2024). Given that CathB6 interacts with MORF2 and recapitulates MORF2-OX and *gun1* transcriptional signatures (Figure 4), we examined whether CathB6 affects MORF2-dependent chloroplast RNA editing.

After filtering sites for sufficient read depth and sequence quality, we determined editing efficiency at 25 sites. Among these, only *clpP*-559 of the *clpP* transcript showed a significant change in editing efficiency between CathB6-GFP and GFP control plants (Binomial GLM, *P* < 10^-7^, Fig. S4A, B). No significant changes were detected between *eds1-2* and wild-type Col-0 plants (Figure S4B). This C-to-T editing event at position 559 results in a histidine-to-tyrosine substitution in the ClpP protein (Figure S4C). The editing of *clpP*-559 is repressed during dark-to-light transition via GUN1 and CONSTITUTIVELY PHOTOMORPHOGENIC 1 (COP1) (Hu et al., 2025), and the reduced editing leads to decreased ClpP-mediated proteolysis, thereby promoting plastid biogenesis (Majeran et al., 2019; Hu et al., 2025). Therefore, these data suggest that CathB6 does not have a major impact on the editing of chloroplast-encoded transcripts, except for *clpP*-559, supporting the idea that CathB6 promotes plastid biogenesis.

### In the presence of GLK2, CathB6 localizes to nuclei and is depleted from p-bodies

To investigate subcellular localization of CathB6 and its interactors, we co-infiltrated CathB6-GFP with GLK2-RFP or MORF2-mCherry into *N. benthamiana* leaves. As expected, GLK2-RFP located within plant cell nuclei in the presence of free GFP (Figure 5A). Moreover, CathB6-GFP associated with cytoplasmic punctate structures, identified as p-bodies, in the cytoplasm in the presence of free RFP or mCherry (Figure 5A), in agreement with previous reports (Liu et al., 2025). However, CathB6-GFP located in nuclei in the presence of GLK2-RFP (Figure 5A), indicating that GLK2 removes CathB6 from p-bodies and relocates this aphid effector to nuclei.

**Figure 5.**
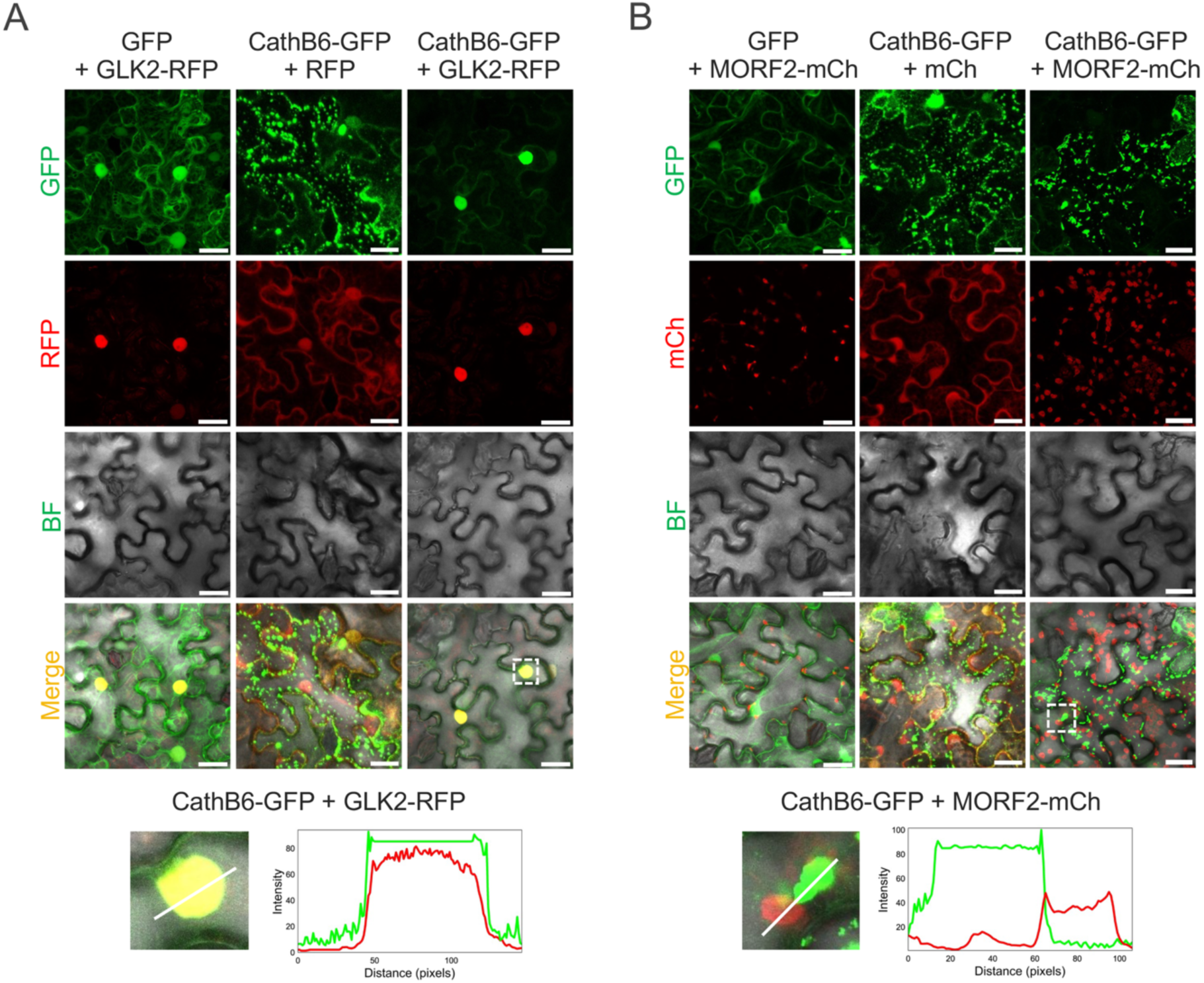
CathB6 localizes to the nucleus in the presence of nuclear GLK2 but remains associated with p-bodies in the presence of chloroplast-localized MORF2. Confocal images showing the localizations of GFP or CathB6-GFP in the presence or absence of GLK2-RFP (A) or MORF2-mCherry (B) in *N. benthamiana* cells. Scale bars, 30 μm. Graphs below the confocal images are intensity profiles along the marked lines of the puncta indicated with white boxes in the merged confocal images above.

MORF2-mCherry localizes to chloroplasts in the presence of the GFP control (Figure S5), in agreement with MORF2 being predominantly located within chloroplasts (Huang et al., 2019). CathB6-GFP remained located in p-bodies in the presence of MORF2-mCherry and there was no obvious localization of MORF2-mCherry into p-bodies. Therefore, MORF2 and CathB6 do not appear to relocate each other. Nonetheless, it is interesting that CathB6-GFP-labeled p-bodies were frequently observed adjacent to, or closely associated with, the chloroplast periphery (Figure 5B; Figure S5). We cannot exclude the possibility that CathB6 association with MORF2 is transient, potentially occurring in the cytoplasm, or in p-bodies, during or shortly after MORF2 is translated. Such an interaction may not be detected given that the majority of MORF2 is in the chloroplast.

### The CathB6 proximal interactome is enriched in proteins linked to chloroplasts, stress responses, RNA metabolism and intracellular transport

To further characterize CathB6-mediated modulation, we determined the CathB6 proximal interactome using TurboID-based proximity labelling in *A. thaliana* Col-0 plants coupled with mass spectrometry (PL-MS). These data were previously released (Liu et al., 2024) and were further analyzed herein. We identified 267 *A. thaliana* proteins significantly enriched in CathB6-TurboID samples compared to GFP-TurboID controls, based on a fold-change threshold (CathB6-TurboID/GFP-TurboID ratio > 2) and a statistical significance threshold (*P* < 0.05 from two-tailed Student’s t-test) (Table S5). GO: Biological Process and GO: Cellular Component analysis showed strong enrichments for chloroplast-related compartments and processes, and GO: Molecular Function in ATP binding, mRNA binding and unfolded protein binding (Table S6).

Further manual annotations revealed more than half of the proteins (54%,144 proteins) being associated or located in chloroplasts and the remainder in the cytosol or other compartments (27%), nuclei (7%), mitochondria (4%) or uncharacterized locations (8 %) (Figure S6A; Table S7). Functional categories comprised chloroplast processes (43%), abiotic and biotic stress responses (25%) and RNA-related processes (12%) (Fig. S6B, C; Table S7). Cross-category analysis showed significant overlap with 28 of 130 chloroplast proteins being linked to stress responses and 7 of 35 RNA-binding proteins to abiotic/biotic processes (Figure S6D).

The CathB6 proximal interactome included EDS1 and Acd28.9 (Table S5), consistent with their direct binding to CathB6 and their opposing effects on p-body localization, with CathB6 recruiting EDS1 and its signalling partners to p-bodies, whereas Acd28.9 depleting them from these compartments. (Liu et al., 2025). This indicates that proximity labeling experiment was succesful. However, the GLKs, MORFs, and GUN1 were not detected. This may be explained by their low abundance (Tadini et al., 2020) and their interactions with CathB6 possibly being transient. Nonetheless, the CathB6 interactome did include other GUN proteins, including GUN4, which directly interacts with MORF2 (Huang et al., 2019; Yuan et al., 2022), as well as GUN4-interacting proteins GUN5, EX1, and EX2 (Li et al., 2022) (Figure 6; Table S5). GUN4 and GUN5 have been reported to localize to p-bodies together with the m^6^A reader ECT1 (Jang et al., 2019), which was also identified in the CathB6 interactome (Table S5) and promotes decay of SA-induced transcripts to attenuate stress responses (Jang et al., 2019; Lee et al., 2024). Additionally, the interactome included the DEAD-box RNA helicase RH50 (Table S5), which colocalizes with GUN1 (Paieri et al., 2018). The CathB6 interactome also included PAP3 (PTAC10), PAP5 (PTAC12), PAP8 (PTAC6), PAP13 (FLN2), PTAC4 and PAP26 (Table S5), which are components of the plastid-encoded RNA polymerase (PEP) complex (Vergara-Cruces et al., 2024; Liu et al., 2026). These findings support the idea that CathB6 modulates chloroplastic processes.

**Figure 6.**
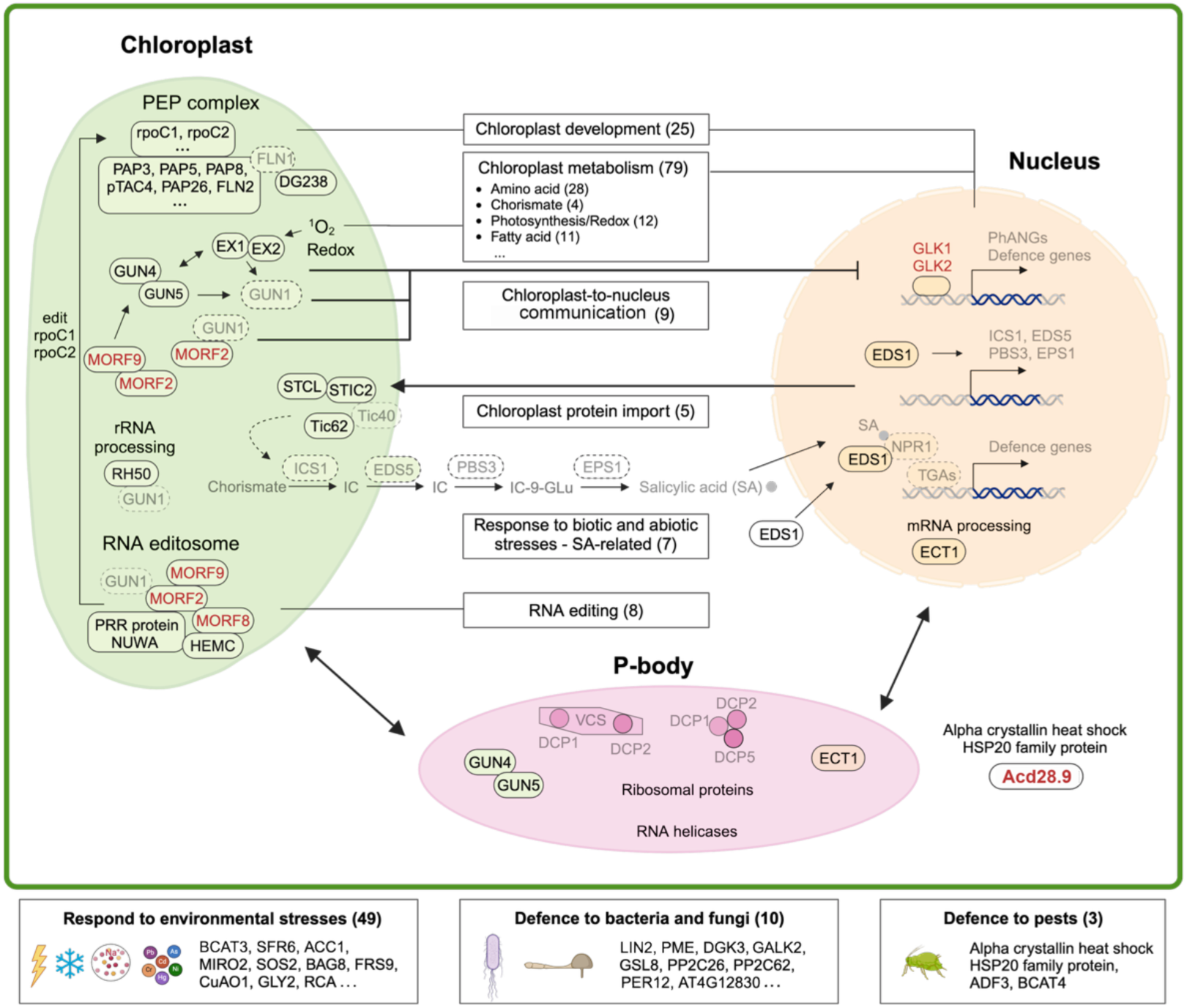
Interacting network of 267 proteins identified by CathB6-TurboID. Proteins identified by PL-MS at a ratio of CathB6-TurboID/GFP-TurboID > 2 are printed in black, those identified as CathB6 interactors in Y2H experiments in red, and those in the same pathways but not detected by TurboID PL-MS in light grey. Specific processes are shown within boxes in the central area of the figure with numbers of proteins involved in these that were identified at CathB6-TurboID/GFP-TurboID > 2 indicated between parentheses. Other proteins identified at a ratio of CathB6-TurboID/GFP-TurboID > 2 involved in plant responses to the environment, defence to bacteria and fungi and defence to pests, including aphids, are shown in boxes at the bottom. PhANGs, photosynthesis-associated nuclear genes.

We previously showed that CathB6-labelled p-bodies are highly mobile (Liu et al., 2025). Consistent with this, the proximal interactome contained proteins involved in actin and tubulin organization and motility, including four Prefoldin (PFD) proteins (PFD2, PFD4, PFD5, and PFD6), three TCP-1/Cpn60 chaperonins (CCT2, CCT6-2, and CCT8), myosin motor XI MYOSIN H (XIH), and actin depolymerizing factor ADF3 (Gu et al., 2008; Peremyslov et al., 2008; Rodríguez-Milla and Salinas, 2009; Fichtenbauer et al., 2012; Ahn et al., 2019; Ding et al., 2022) (Figure 6). Notably, ADF3 is also required for Arabidopsis defense to *M. persicae* (Mondal et al., 2018).

Thus, the CathB6 proximity interactome is dominated by proteins involved in chloroplast functions, (a)biotic interactions, RNA metabolism and intracellular transport cytoskeletal components.

## DISCUSSION

We previously found that the *M. persicae* effector CathB6 increases *A. thaliana* susceptibility to the aphid and sequesters the central plant immune regulatory protein EDS1 and its signalling partners, PAD4 and ADR1, to p-bodies (Liu et al., 2025). Here, we found that there is substantial overlap in up- and downregulated transcripts in the same directions between CathB6 and *eds1-2* lines, indicating that CathB6 suppresses EDS1-controlled processes. Nonetheless, CathB6 transgenic lines also displayed transcriptional reprogramming that did not overlap with those of *eds1-2* lines. Thus, although EDS1 is an important CathB6 target, CathB6 appeared to affect additional host pathways that contribute to aphid susceptibility.

A yeast two-hybrid screen of CathB6 against a cDNA library of *A. thaliana* exposed to *M. persicae*, other insects and a pathogen, and follow-up experiments, identified GLK1, GLK2 and several MORF proteins as additional CathB6 interactors as well as Acd28.9, which was previously identified as an interactor of aphid CathB proteins (Liu et al., 2025).

CathB6 also appears to influence the expression of transcripts regulated by GLK1 and GLK2, which are GARP-family transcription factors that act as master regulators of PhANGs (Fitter et al., 2002; Waters et al., 2009; Chen et al., 2016; Li et al., 2022; Zheng et al., 2024). Transcripts differentially regulated in CathB6 lines overlapped with those altered in the *glk1.1 glk2.1* mutant, although the majority showed opposite patterns of regulation with predominant upregulation of genes in CathB6 and downregulation in *glk1.1 glk2.1*. This suggests that CathB6 is unlikely to inhibit GLK function. Instead, CathB6 may promote GLK transcriptional activity and/or, given its association with p-bodies, influence the stability of GLK-regulated transcripts. This hypothesis is in agreement with aphid fecundity assays, as we found that aphids produced more progeny on lines stably expressing CathB6, confirming previous results (Liu et al., 2025), whereas aphid fecundity was reduced on *glk1.1 glk2.1* plants. In addition, CathB3 and CathB6, which both promote aphid fecundity (Liu et al., 2025) interact with the GLKs, whereas CathB9 does not promote aphid fecundity (Liu et al., 2025) and does not interact with GLKs. This indicates that GLK activity is important for aphid performance.

The link between CathB6, GLKs and chloroplast-to-nucleus signalling is further supported by the proximity of CathB6 to GUN4, GUN5, EX1 and EX2. GUN4 and GUN5 are key components in tetrapyrrole metabolism and chlorophyll biosynthesis (Waters et al., 2009; Kim et al., 2023). Tetrapyrrole intermediates, either free or associated with the GUN4–GUN5 complex, contribute to chloroplast-to-nucleus signalling that suppresses GLK activity (Larkin, 2016). GUN1 also interacts with MORF2 to suppress GLK-mediated PhANG expression following chloroplast damage (Koussevitzky et al., 2007; Tokumaru et al., 2017). GUN proteins form complexes with EX1 and EX2 (Dogra et al., 2022; Li et al., 2023). EX1 mediates singlet oxygen-induced stress signalling, whereas EX2 attenuates EX1 sensitivity (Wagner et al., 2004; Dogra et al., 2022). Consistent with a role in this pathway, genes downregulated in CathB6-GFP versus GFP lines were enriched for responses to oxygen levels, hypoxia, stress and stimuli. Together with the strong overlap between the CathB6 and *gun1-9* transcriptomes, these findings suggest that CathB6 interferes with the GUN- associated chloroplast-to-nucleus signalling axis to downregulate oxygen-induced stress levels and sustain chloroplast activity.

Within the MORF protein family, CathB6 shows the strongest interactions with MORF2, MORF3, MORF6, MORF8 and MORF9, no interaction with MORF1 and weaker interactions with other MORF proteins. This specificity suggests that CathB6 does not bind MORFs indiscriminately. Transcriptomic comparisons further support a functional link between CathB6 and MORF-associated processes, as differentially expressed genes in the CathB6 transgenic plants overlapped with those in NF-treated MORF2-OX relative to NF-treated wild-type controls, such as the enhanced expression of PhANGs (Zhao et al., 2019), with most changing in the same direction. The decreased editing of the *clpP* transcript also suggest that CathB6 promotes plastid biogenesis (Majeran et al., 2019; Hu et al., 2025).

Although MORFs predominantly function in chloroplasts, CathB6 was not detected within this organelle. Instead, CathB6 remained localized to cytoplasmic p-bodies in the presence of MORF2. Because these chloroplast-associated proteins (Yuan et al., 2022) are nuclear-encoded, translated in the cytosol and likely imported into chloroplasts via the TIC/TOC machinery, CathB6 may interact with MORF-containing or plastid precursor complexes before chloroplast import. This model is consistent with the enrichment of PhANG-encoded proteins in the CathB6 proximal interactome and with reports that *gun1* mutants have reduced plastid protein import capacity, leading to retention of plastid precursor proteins in the cytosol (Tadini et al., 2020). However, aphid fecundity was slightly increased on the *morf2-1* mutants and CathB9, which does not promote aphid fecundity (Liu et al., 2025), also binds MORF proteins. It should also be taken onto account that the MORF2-OX lines can also produce a weak *morf2* loss-of-function-like phenotype (Zhao et al., 2019). Therefore, CathB-mediated modulation of MORF proteins may be more complicated and need further investigation.

The CathB6 interactome was also enriched for proteins involved in actin/tubulin biogenesis, cytoskeletal organization and actin-based motility, including prefoldin complex components, ADF3 and myosin XIH. Notably, ADF3 is required for Arabidopsis defense against *M. persicae* and for PAD4 induction (Mondal et al., 2018). Because myosin XI members bind the p-body component DCP1 and regulates p-body movement (Steffens et al., 2014), CathB6 may co-opt cytoskeletal trafficking to alter the positioning or sequestration of defense-related mRNA-protein complexes. In this context, it is interesting that *M. persicae* fecundity increase on *A. thaliana eds1* mutant plants is more moderate than on *A. thaliana pad4* and triple *adr1* mutant plants (Liu et al., 2025) and that the PAD4 lipase domain can promote *A. thaliana* resistance to *M. persicae* on its own (Dongus et al., 2019). *M. persicae* CathB effectors may suppress EDS1 highly effectively while allowing its signalling partners, PAD4 and ADR1, to retain partial activity in a process that involves ADF3.

In conclusion, our findings support a model in which aphid CathB effectors suppress plant stress responses while sustaining chloroplast activity (Figure 7). During feeding, aphids deliver CathB effectors into plant cells, where CathB6 localizes to p-bodies and recruits EDS1, together with PAD4 and ADR1, to these compartments (Liu et al., 2025). This recruitment is associated with the suppression of plant immune responses, including EDS1-regulated immunity (Liu et al., 2025; this study). CathB effectors also interact with GLK transcription factors, which promote the relocalization of CathB6 to the nucleus and reduce its association with p-bodies. In parallel, CathB6 modulates the GLK–MORF2–GUN1 signalling pathway in a manner consistent with enhanced GLK and MORF2 activity and reduced GUN1 activity. CathB effectors also interact with selected MORF proteins, including MORF2, although whether these interactions occur in planta remains unknown. These activities may benefit aphids by simultaneously suppressing EDS1-dependent defences and maintaining chloroplast functions that support primary metabolism and nutrient production. Thus, rather than simply disabling host immunity, CathB6 reconfigures plant cellular pathways to establish a more permissive and nutritionally favourable environment for aphid growth and reproduction.

**Figure 7.**
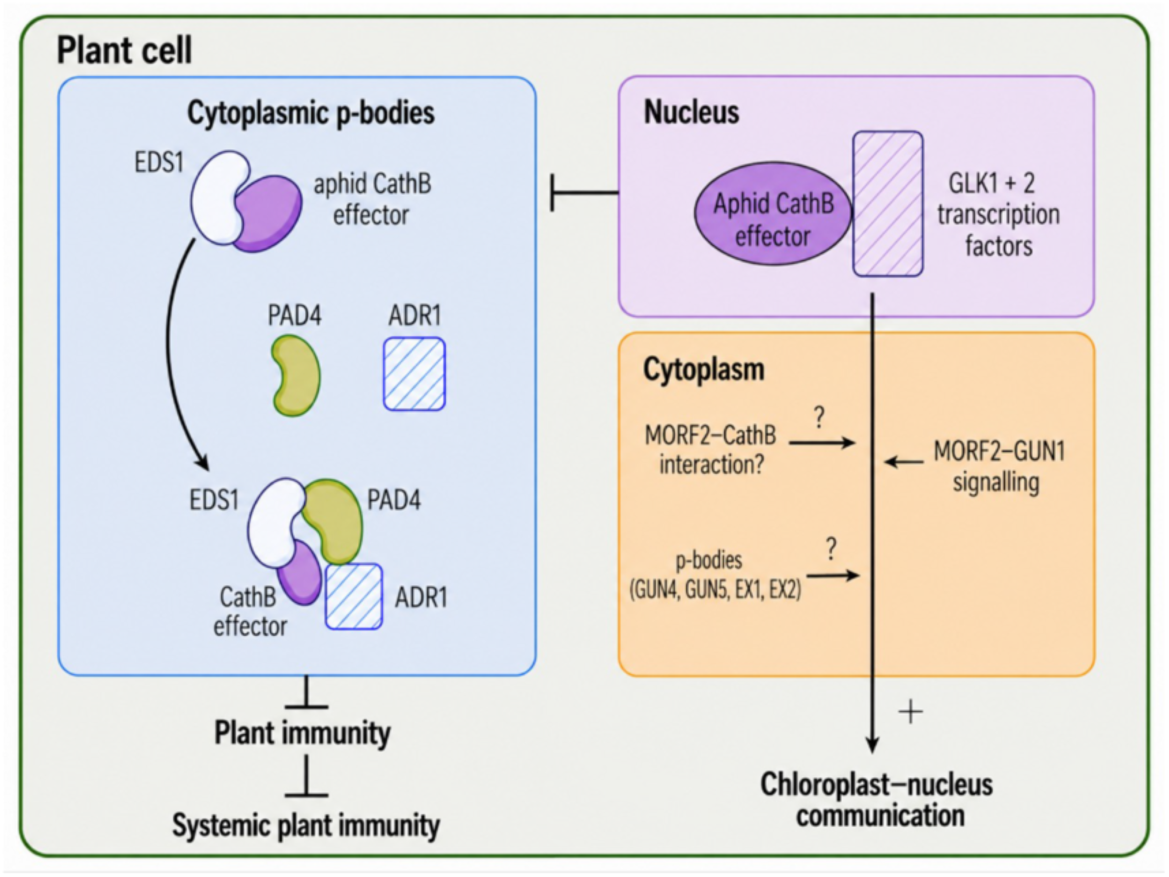
Model for *M. persicae* CathB effector function in modulating MORF–GUN– GLK signalling to maintain cellular homeostasis during aphid feeding. During feeding, aphids secrete CathB effectors into plant cells. Once inside the host cell, CathB exhibits two distinct activities. Left panel: CathB6 localizes to processing bodies (p-bodies). CathB6 also interacts with EDS1 and recruit EDS1, together with its partners PAD4 and ADR1, to the p-bodies (Liu et al., 2025). CathB6 suppresses plant immune responses (Liu et al., 2025), including EDS1-regulated immunity, as shown in this study. Right panel: CathB effectors interact with plant GLK transcription factors, which recruit CathB6 to the nucleus and reduce its localization to p-bodies. CathB6 modulates the GLK–MORF2–GUN1 pathway in a manner that maintains chloroplast– nucleus communication. Aphid CathB effectors also interact with specific members of the MORF family, including MORF2, although whether this interaction occurs *in planta* remains unclear. The CathB6 proximal interactome further links p-body dynamics, nucleus-chloroplast communications, and cytoskeletal organization with defence-related pathways. Together with evidence that p-bodies contribute to nucleus-chloroplast communications and contain GUN4 and GUN5 (Jang et al., 2019), these findings suggest that CathB6 may coordinate p-body-associated processes with nucleus-chloroplast communications. However, how CathB modulates these pathways remains unresolved, as indicated by the question marks.

## MATERIALS AND METHODS

### Aphid colony maintenance

The *M. persicae* Clone O was reared on *A. thaliana* Col-0 and maintained in a growth chamber (20°C, 14 h light / 10 h dark, 75% humidity) since 2010 (Mathers et al., 2017).

### Plant lines and growth conditions

All *A. thaliana* lines used in this study are listed in Table S8. Stable transgenic *A. thaliana* Col-0 lines expressing GFP and CathB6-GFP (*M. persicae* CathB6 gene ID: MYZPE13164_O_EIv2.1_0111090, Liu et al., 2024) were generated previously (Liu et al., 2025). From these, Col-0_GFP (lines 3-1 and 3-5) and Col-0_CathB6-GFP (lines 2-7 and 4-5) were used for RNA-seq analyses in this study, and lines Col-0_GFP_3-5 and Col-0_CathB6-GFP_4-5 were used for aphid fecundity assays. The Col-0 *eds1-2* mutant was generated by introgressing the Ler-0 *eds1-2* null allele into the Col-0 background through an initial cross followed by eight successive backcrosses (Bartsch et al., 2006). Lines expressing GFP-TurboID or CathB6-TurboID for proximal interactome experiments have been described (Liu et al., 2024). The *A. thaliana* mutants *morf2-1*, *glk1.1 glk2.1* and *acd28.9* were obtained from the NASC Arabidopsis stock centre.

Wild-type, mutant or transgenic *A. thaliana* Col-0 plants used for fecundity assays were maintained under short day conditions (10 h light/14 h dark) at 22°C with a humidity of 70%. *N. benthamiana* plants for confocal microscopy were grown under long day photoperiod (16 h light/8 h dark) at 22°C with a humidity of 80%.

### RNA-seq analyses

Seeds of wild-type Col-0, transgenic *A. thaliana* lines expressing GFP or CathB6- GFP, and the *eds1-2* mutant were sown on MS plates and grown to ten-day-old seedlings. RNA was extracted from three replicate samples of pooled seedlings for two independent lines each of CathB6-GFP and GFP empty vector controls, and from *eds1-2* mutants and wild-type Col-0 control. Directional library prep and sequencing was performed by Novogene (Cambridge, UK), yielding approximately 30 million reads per sample. FastQC (Andrews, 2010) and Trim Galore (Krueger et al., 2023) were used to quality control data and trim reads. Reads were mapped to the *Arabidopsis thaliana* TAIR10 annotation using HISAT2 (Kim et al., 2019), and transcripts quantified using HTSeq-count (Putri et al., 2022). Principal component analysis was performed and Differentially expressed genes were determined using DESeq2 (Love et al., 2014). Venn diagrams were plotted with Venny 2.1 online software (https://bioinfogp.cnb.csic.es/tools/venny/index.html), and Gene Ontology enrichment analysis was performed using Panther (Mi et al., 2013). RNA sequence reads are available from the NCBI Sequence Read Archive with project number PRJNA1431337. Comparisons to published transcriptomes were done using DE gene lists provided for the *glk1.1 glk2.1* mutant (Ahmad et al., 2019) and the MORF2-overexpressor and *gun1-9* mutant (Zhao et al., 2019).

### RNA editing analyses

Chloroplast genome-encoded transcripts were analysed for editing efficiency using Freebayes 1.2.0 (Garrison and Marth, 2012). BAM files generated from HISAT2 were used to call variants from the chloroplast genome, and resulting VCF files were filtered to remove sites with average coverage <10 across all samples. Identified editing sites covered 25 out of 40 previously described sites (Castandet et al., 2016). Percentage editing efficiency relative to the control plants (GFP or wild-type Col-0) was calculated, and significant changes in editing efficiency was determined using a Binomial GLM.

### GO ontology enrichment analysis

Functional ontology (GO) analyses of identified proteins were conducted on DAVID (https://david.ncifcrf.gov/summary.jsp) and plotted with R package ggplot2 (Wickham, 2011). GO annotation of Biological Process (BP), Cellular Component (CC) and Molecular Function (MF) were analyzed with FDR < 0.05.

### Plasmids

All plasmids used are listed in Table S9 along with references. The constructions of plasmids that were generated in this study are described below.

### Yeast two-hybrid (Y2H) library screening

The coding sequence corresponding to the mature domain of CathB6 (residues Arg61-Asn338; *M. persicae* CathB6 gene ID: MYZPE13164_O_EIv2.1_0111090, Liu et al., 2024) was amplified from *M. persicae* clone O cDNA and cloned into the pLexA-N bait vector. The resulting pLexA-N-CathB6 construct was transformed to *E. coli*, and the plasmid was verified by Sanger sequencing. A yeast-2-hybrid library was constructed using combined mRNAs isolated from non-infected *Arabidopsis thaliana* Col-0 plants and *A. thaliana* Col-0 plants exposed to the aphid species *Myzus persicae* (peach-potato aphid) or *Acyrthosiphon pisum* (pea aphid), leafhopper species *Macrosteles quadrilineatus* (aster leafhopper) or *Dalbulus maidis* (corn leafhopper), or *A. thaliana* Col-0 plants infected with the Aster Yellows Witches Broom (AY-WB) phytoplasma (*Candidatus* Phytoplasma asteris) (Gravino et al., 2024). The cDNAs were cloned into the SfiI site of pGAD-HA prey vector (Dualsystems).

For library screening, the pLexA-N-CathB6 bait construct was transformed into the yeast strain NMY51, and transformants were selected on SD-trp plates at 28 °C for 5 days. A single yeast colony expressing pLexA-N-CathB6 was inoculated into liquid SD-trp culture and grown overnight at 28 °C with shaking. The next day, yeast cells were harvested and made competent using the PEG/LiOAc method. The pGAD-HA library plasmids were then transformed into the competent yeast cells expressing the bait construct at a serial dilutions of 1:100, 1:1000 and 1:10000. Transformants were screened on a series of selective media, including triple dropout medium lacking leucine, tryptophan, and histidine (SD-LWH) supplemented with 5 mM or 10 mM 3- amino-1,2,4-triazole (3-AT), and quadruple dropout medium lacking leucine, tryptophan, histidine, and adenine (SD-LWHA).

Positive colonies were picked and inoculated into liquid medium under the same selective conditions. Plasmids were extracted from yeast and re-transformed to *E. coli*. Positive *E. coli* colonies were inoculated, and plasmid DNA was extracted and subjected to Sanger sequencing. The resulting sequences were analyzed by BLAST against the TAIR database (https://www.arabidopsis.org/tools/blast/) to identify candidate *A. thaliana* genes.

### Y2H assay

To validate the Y2H library screening results, Y2H assays were performed using two yeast strains, NMY51 and AH109, with strain-specific vector systems. For the NMY51 strain, the coding sequences of candidate genes were amplified and cloned into pGAD-HA (AD) vector. These constructs were then co-transformed with pLexA-N-CathB6 (BD) construct. Empty pGAD-HA and pLexA-N vectors were used as negative controls. For the AH109 strain, candidate gene coding sequences were amplified and cloned into the pDEST-GADT7 (AD) vector, while the sequence encoding the mature domain of CathB6 (residues Arg61-Asn338) was cloned into pDEST-GBKT7 (BD) vector. The resulting AD and BD constructs were co-transformed into AH109, with empty pDEST-GADT7 and pDEST-GBKT7 vectors serving as negative controls.

For additional interaction tests, the assay was conducted in AH109 strain. The full length coding sequences of *A. thaliana* GLK1 (AT2G20570), GLK2 (AT5G44190) and MORF1 through MORF9 proteins (AT4G20020, AT2G33430, AT3G06790, AT5G44780, AT1G32580, AT2G35240, AT1G72530, AT3G15000 and AT1G11430, resepctively) were amplified and constructed into the pDEST-GADT7 vector, and co-transformed with pDEST-GBKT7-CathB6. Those carrying sequences of *A. thaliana* EDS1 (AT3G48090) and Acd28.9 (AT5G47590) were constructed earlier (Liu et al., 2025).

Transformants were first assessed on solid double dropout medium lacking leucine and tryptophan (SD-LW) to confirm the presence of both AD and BD constructs. Protein-protein interactions were subsequently assessed on triple dropout medium lacking leucine, tryptophan, and histidine (SD-LWH) supplemented with 5 mM 3- amino-1,2,4-triazole (3-AT), and on quadruple dropout medium lacking leucine, tryptophan, histidine, and adenine (SD-LWHA). All plates were incubated at 28 °C for 5 days prior to imaging. Each transformation combination was repeated at least three times.

### Coimmunoprecipitation (Co-IP) in *N. benthamiana*

The coding sequences corresponding to the mature domain of *M. persicae* CathB6 (residues Arg61-Asn338) was cloned into the pB7FWG2 vector. The full-length coding sequences of *A. thaliana* GLK1, GLK2, MORF1 and MORF2 were amplified, and N-terminally tagged with 3×HA, and ligated into the Gateway vector pB7WG2. All constructs were separately transformed into *A. tumefaciens* strain GV3101 and co-infiltrated into leaves of 4-week-old *N. benthamiana* plants. Infiltrated leaves were harvested between 48 and 72 hours post-infilatration (hpi).

Total proteins were extracted using extraction buffer [150 mM Tris-HCl (pH 7.5), 150 mM NaCl, 10 mM EDTA, 10% Glycerol, 20 µM NaF, 10 mM DTT, 0.5% (w/v) PVPP, 1% protease Inhibitor cocktail (Sigma), 0.2% Igepal]. GFPtrap beads (Chromotech) were added to the protein extracts and incubated on a rotating wheel at 4 °C overnight. The next day, the beads were washed six times with washing buffer [10 mM Tris-HCl (pH 7.5), 150 mM NaCl, 0.5 mM EDTA, 0.2% Igepal]. After washing, proteins were eluted from the beads using 4 × LDS Sample Loading Buffer containing 10 mM DTT, separated by 12% SDS-PAGE (Invitrogen), and transferred onto 0.45 μm PVDF membranes. The membranes were then probed with anti-GFP (Santa Cruz) and anti-HA (BioLegend) antibodies.

### Aphid fecundity assay

Seeds from wild-type Col-0, *eds1-2*, *morf2-1*, *glk1.1 glk2.1*, and *acd28.9* mutant *A. thaliana* lines (Table S8) were sown on MS medium. Homozygous GFP or CathB6- GFP transgenic lines (Table S8) were sown on MS medium supplemented with BASTA to confirm the presence of the transgene. Seedlings were transplanted to individual pots and grown in a controlled environmental room (CER) at 22°C under a 10 h light/14 h dark photoperiod.

Prior to the assay, adult *M. persicae* (Clone O) were transferred from the stock colony to three-week-old *A. thaliana* plants and enclosed in sealed cages. The next day, newly deposited nymphs were transferred to test plants at one nymph per plant. Nymphs matured and began reproducing after one week; progeny were counted on days 7, 9, 11, 13, and 15 post-transfer. Fecundity was calculated as the number of nymphs produced per adult. Each line was initially tested with 15 ramdomly selected plants; due to occasional aphid loss during development, final replicates ranged from 10 to 15 per line.

Statistical analysis was performed using one-way ANOVA followed by Tukey’s multiple comparison test (GraphPad Prism 11), with *P* < 0.05 considered significant. Boxplots were generated using R (v4.6.0).

### Subcellular localization

The coding sequence corresponding to the mature domain of *M. persicae* CathB6 (residues Arg61-Asn338) was cloned into the pB7FWG2 vector for GFP tagging. The full-length coding sequences of GLK2 and MORF2 were cloned into pB7RWG2 (RFP tag) and pB7WG2-mCherry (mCherry tag), respectively, using Gibson assembly and Gateway cloning.

The resulting constructs were transformed into *A. tumefaciens* strain GV3101, plated on LB solid medium containing appropriate antibiotics, and incubated at 28 °C for 24-48 hours. Colonies were picked and verified by PCR with gene-specific primers. Positive colonies were cultured overnight at 28 °C, harvested by centrifugation, and resuspended in infiltration buffer (10 mM MgCl₂, 10 mM MES, pH 5.6) supplemented with 100 μM acetosyringone.

For leaf infiltration, *Agrobacterium* strains expressing GFP or CathB6-GFP were mixed with those expressing RFP, GLK2-RFP or MORF2-mCherry, along with the silencing suppressor pCB301-P19, each adjusted to an OD_600_ of 0.3. The mixtures were infiltrated into the abaxial surface of leaves from 4-week-old *N. benthamiana* plants using a 1 mL needleless syringe. Infiltrations were performed on randomly selected leaves from two independent plants. Leaf samples were collected for imaging at 48-72 hours post-infiltration (hpi).

For microscopy, live leaf sections were mounted in water. Observations were conducted using a Leica TCS SP8X upright confocal laser scanning microscope equipped with either a 20×/0.75 dry objective or a 63×/1.20 water immersion objective (HC PL APO CS). Sequential, unidirectional scans were performed using the following laser lines: 488 nm (from 65 mW Argon ion laser) for GFP excitation, and 580 nm (from a pulsed SuperK EXTREME supercontinuum white light laser, 470-670 nm, 1.5 mW per line) for RFP/mCherry excitation. Fluorescence emissions were collected at 505-540 nm and 595-620 nm. Images were acquired on hybrid detectors with laser power below 5%, line averaging set to 2, and a pinhole of 1 Airy unit. Gain and z-step settings were adjusted per image. Pixel size was set to 0.18 µm × 0.18 µm, with a pixel dwell time of 600 ns. Each experiment was independently repeated at least three times.

### TurboID-based proximity labelling

TurboID-based proximity labelling coupled with mass spectrometry (PL-MS) to determine the CathB6 proximal interactome in *A. thaliana* Col-0 plants has been previously described (Liu et al., 2024). Analysis of CathB6 PL-MS dataset (Liu et al., 2024) and generation of different plots, including piechart, Venn diagram and barchart, were conducted with R (v4.3.2).

## ACKNOWLEDGEMENTS

We thank the John Innes Centre Entomology, Informatics (and in particular Dan Smith for useful discussions), Bioimaging, Proteomics, and Horticultural Services Technology Platforms for support and advice, as well as members of Hogenhout Lab for useful discussions. This work was funded by UK Research and Innovation (UKRI) Biotechnology and Biological Sciences Research Council (BBSRC) grants to S.A.H. (BB/V008544/1 and BB/R009481/1), with additional support from The Gatsby Charitable Foundation and The Sainsbury Laboratory, Norwich, UK. Further support was provided by the BBSRC Institute Strategy Programmes (BBS/E/J/000PR9797 and BBS/E/JI/230001B) awarded to the John Innes Centre (JIC). The JIC is grant-aided by the John Innes Foundation.

The authors used ChatGPT to assist with grammar and clarity during manuscript editing. All text was reviewed, revised, and verified by the authors, who take full responsibility for the content.

## AUTHOR CONTRIBUTIONS

QL: Conceptualization, Methodology, software, formal analysis, investigation, data curation, writing – original draft preparation, writing – review and editing, visualization. SM: Methodology, software, formal analysis, investigation, data curation, writing – review and editing, visualization. JH: Methodology, software. AN: Methodology, software, formal analysis, investigation. SH: Conceptualization, resources, supervision, methodology, data curation, writing – original draft preparation, writing – review and editing, visualization. All authors have read and agreed tothe published version of the manuscript.

## CONFLICT OF INTEREST

The authors declare that they have no competing interests.

## DATA AVAILABILITY STATEMENT

RNA-seq data are have been deposited in the NCBI SRA project number PRJNA1431337. Mass spectrometry data from the proximity-labelling experiment is described in Liu et al. (2024) and raw data is deposited to the ProteomeXchange Consortium via the PRIDE partner repository with the dataset identifier PXD057789 and 10.6019/PXD057789. All other data supporting the findings of this study are available within the article and its Supplementary Information files.

## Supplementary Materials for

**Figure S1.**
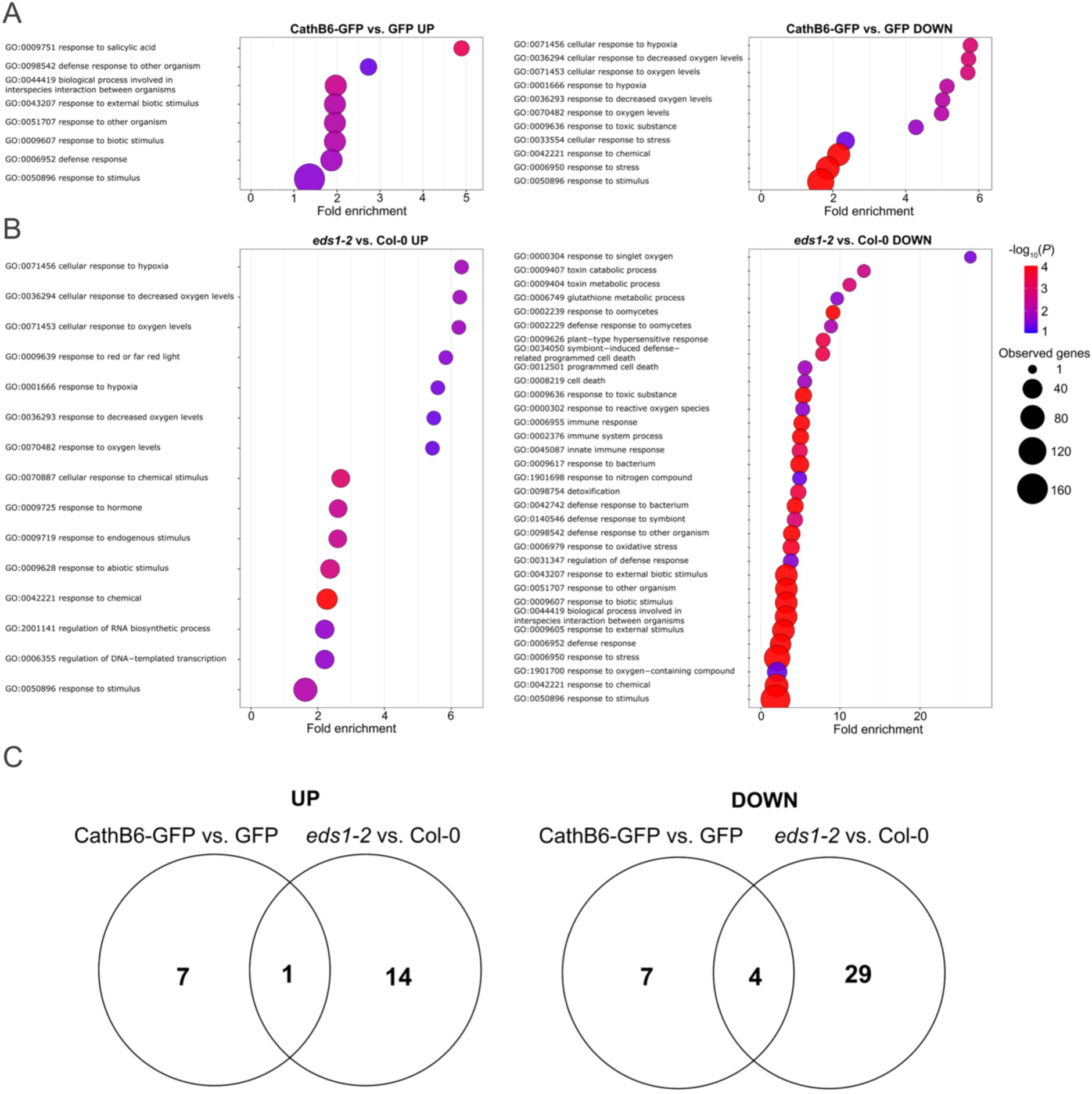
Gene ontology (GO) enrichment analyses of differentially expressed (DE) genes in *A. thaliana* CathB6-GFP transgenic plants and *eds1-2* mutants. (A) Enriched GO terms among DE genes identified in the CathB6-GFP vs. GFP genotypes, up- (left) or down-regulated (right). (B) Enriched GO terms among DE genes identified in the *eds1-2* mutant vs. wild-type Col-0, up- (left) or down-regulated (right). For (A) and (B), the x-axis represents fold enrichment, the dot sizes the number of genes annotated to the term, and the colour gradient the significance level (-log_10_*P* value), ranging from red (most significant) to blue (less significant). (C) Venn diagram showing the overlap of significantly enriched GO terms between CathB6-GFP and *eds1-2* DE genes.

**Figure S2.**
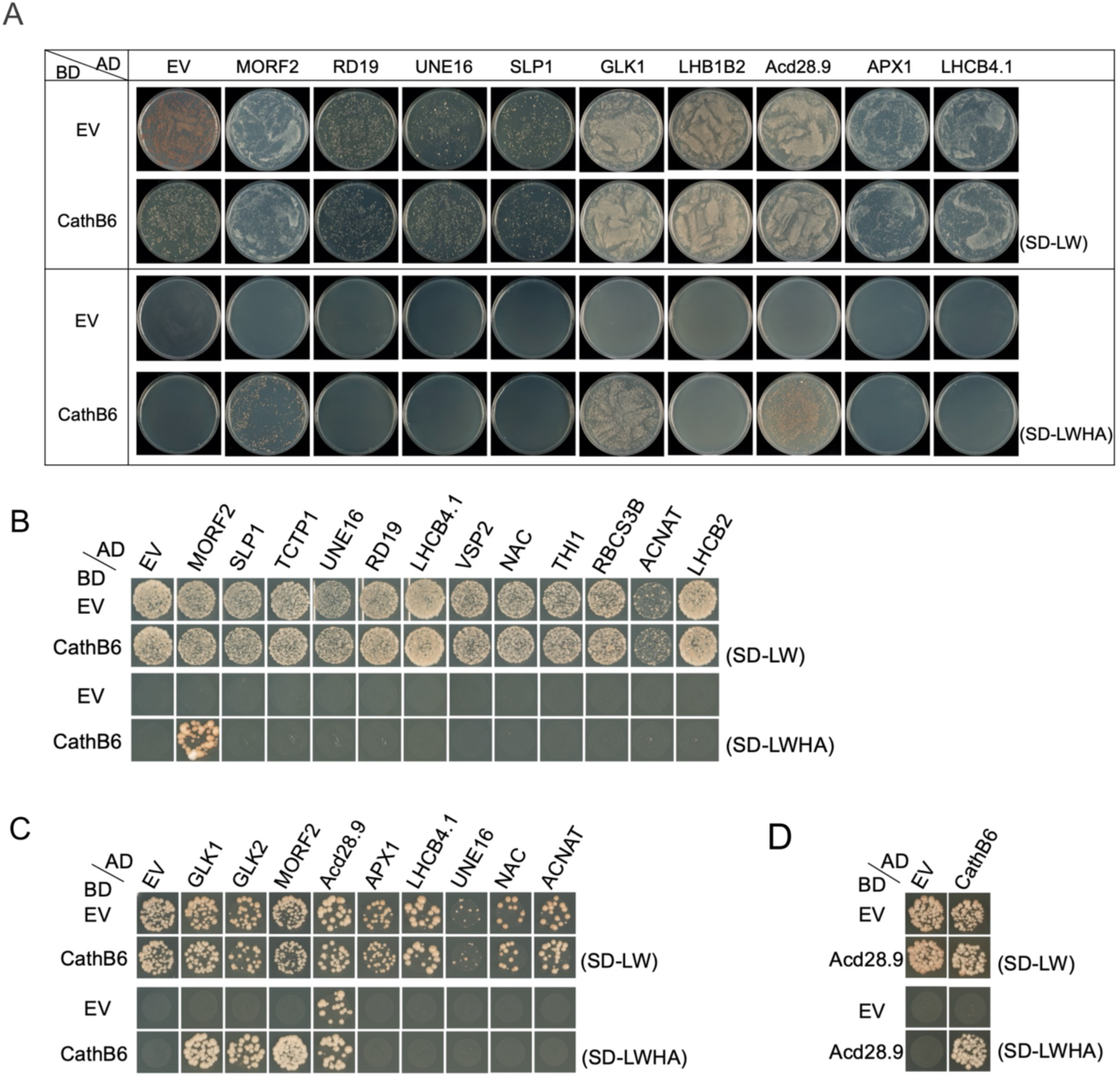
Confirmation of *M. persicae* CathB6 interaction with *A. thaliana* GLK1, MORF2 and Acd28.9. These and the other interactors were identified in a Y2H screen against a cDNA library of *A. thaliana* colonized by *M. persicae*, other insects and a phytoplasma (Supplementary Table 4). Y2H assays were conducted in the yeast NMY51 (A, B) or AH109 (C, D). Abbreviations: EV, empty vector; MORF2, Multiple organellar RNA editing factor 2 (AT2G33430); RD19, Responsive to dehydration 19 (AT4G39090); UNE16, Unfertilized embryo sac 16 (AT4G13640); SLP1, Shewanella-like protein phosphatase 1 (AT1G07010); GLK1, Golden-like 1 (AT2G20570); LHCB1B2, Light harvesting complex photosystem II protein B1B2 (AT2G34420); Acd28.9, Heat shock protein HSP20/alpha crystallin family 28.9 kDa protein (AT5G47590); APX1, Ascorbate peroxidase 1 (AT1G07890); LHCB4.1, Light harvesting complex photosystem II protein 4.1 (AT5G01530); TCTP1, Translationally-controlled tumor protein 1 (AT3G16640); Vegetative storage protein 2, (AT5G24770); NAC, No Apical Meristem domain transcriptional regulator protein (AT3G12910); THI1, Thiazole biosynthetic enzyme (AT5G54770); RBCS3B, Ribulose bisphosphate carboxylase small subunit 3B (AT5G38410); ACNAT, Acyl-CoA N-acyltransferase with RING/FYVE/PHD-type zinc finger domain-containing protein (AT2G37520); LHCB2.1, Light harvesting complex photosystem II protein 2.1 (AT2G05100); AD, GAL4-activation domain; BD, GAL4-DNA binding domain; SD-LW, double dropout medium lacking leucine and tryptophan; SD-LWHA, quadruple dropout medium lacking leucine, tryptophan, histidine, and adenine.

**Figure S3.**
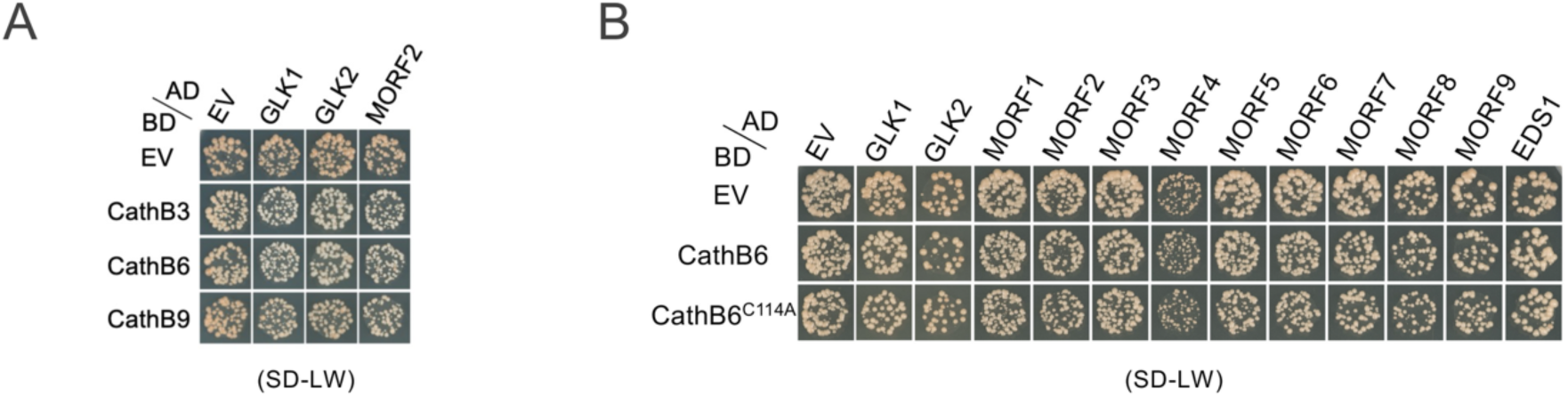
Yeast growth on double dropout medium lacking leucine and tryptophan indicating the expression of AD and BD constructs in Y2H assays shown in Figure 2A (A) and Figure 2C (B) of the main text.

**Figure S4.**
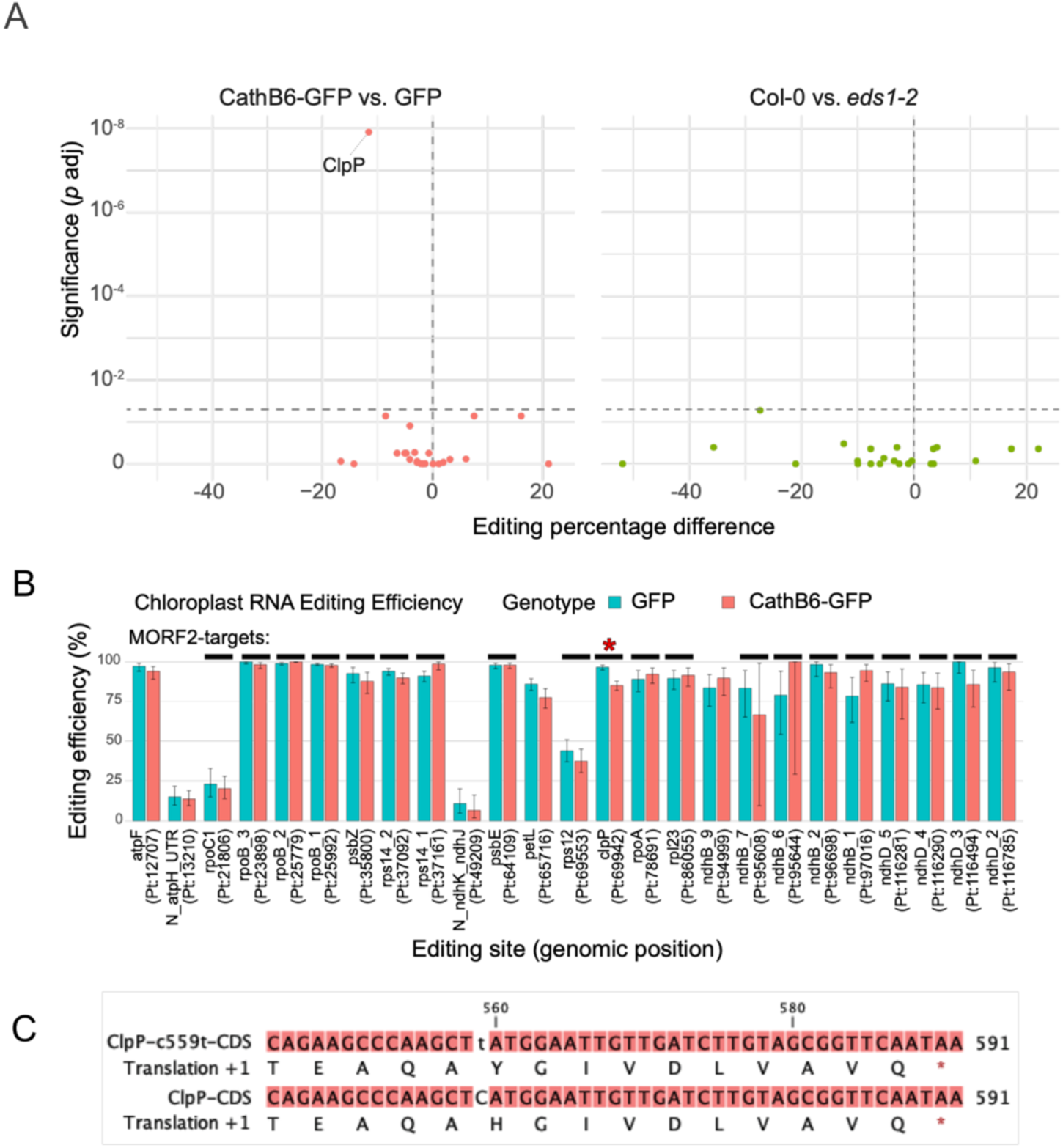
CathB6 reduces the editing efficiency of the chloroplast *clpP* transcript. Transcript editing efficiency at known editing sites was determined using SNP calls from chloroplast genome-mapping RNA-seq reads in CathB6-GFP vs. GFP plants, and *eds1-2* vs. wild-type Col-0 plants. Significant differences in editing efficiency were assessed using a binomial generalized linear model (GLM). Sites were filtered based on read depth, allowing quantification of editing efficiency at 25 of the 40 known sites. (A) Plots of relative editing efficiency against statistical significance in CathB6-GFP vs. GFP (left), and *eds1-2* vs. wild-type Col-0 (right) - only the *clpP* transcript showed a significant reduction in editing efficiency in CathB6-GFP vs. GFP. (B) Absolute editing efficiency at each of the 25 analysed sites. Black bars indicate sites where editing efficiency is altered by MORF2 overexpression, including *clpP*-559 (Zhao et al. 2019). (C) Schematic representation of the C-to-T processing event at position 559 of the *clpP* transcript, resulting in a histidine-to-tyrosine substitution in the encoded ClpP protein.

**Figure S5.**
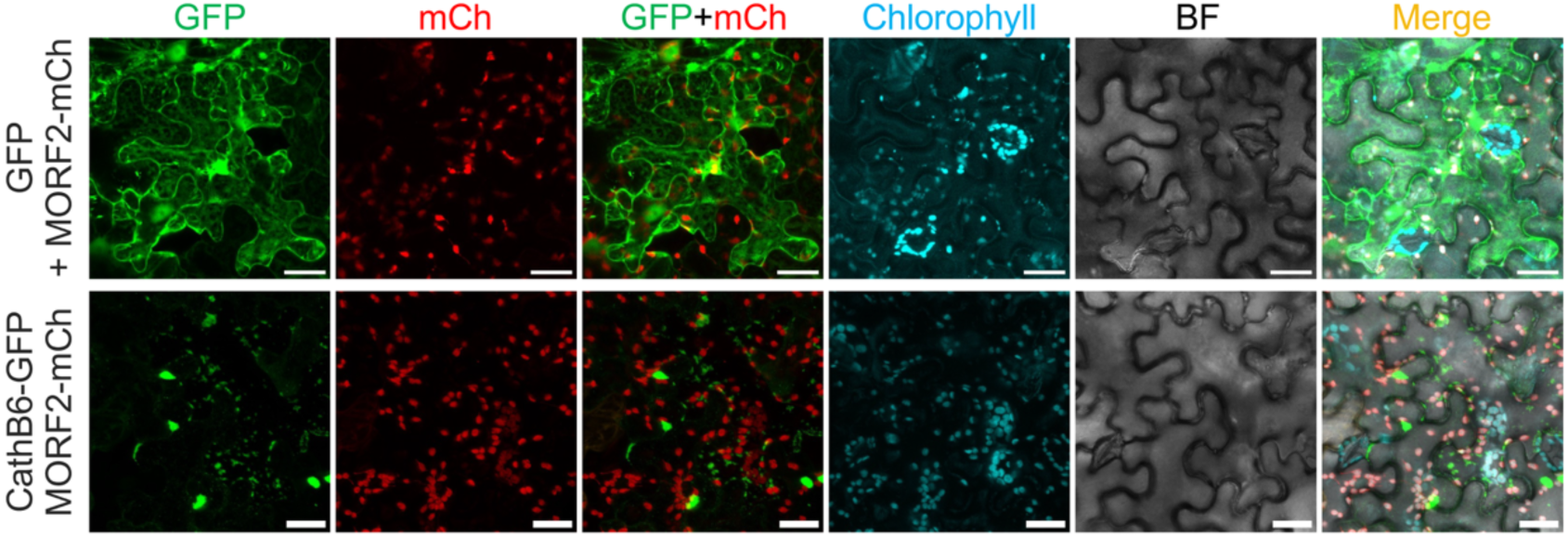
CathB6 does not colocalize with MORF2-labelled chloroplasts in *N. benthamiana* epidermal cells. Confocal images of GFP or CathB6-GFP and MORF2-mCherry. Images were observed with GFP, mCherry or Chlorophyll autofluorescence. Scale bars, 30 μm.

**Figure S6.**
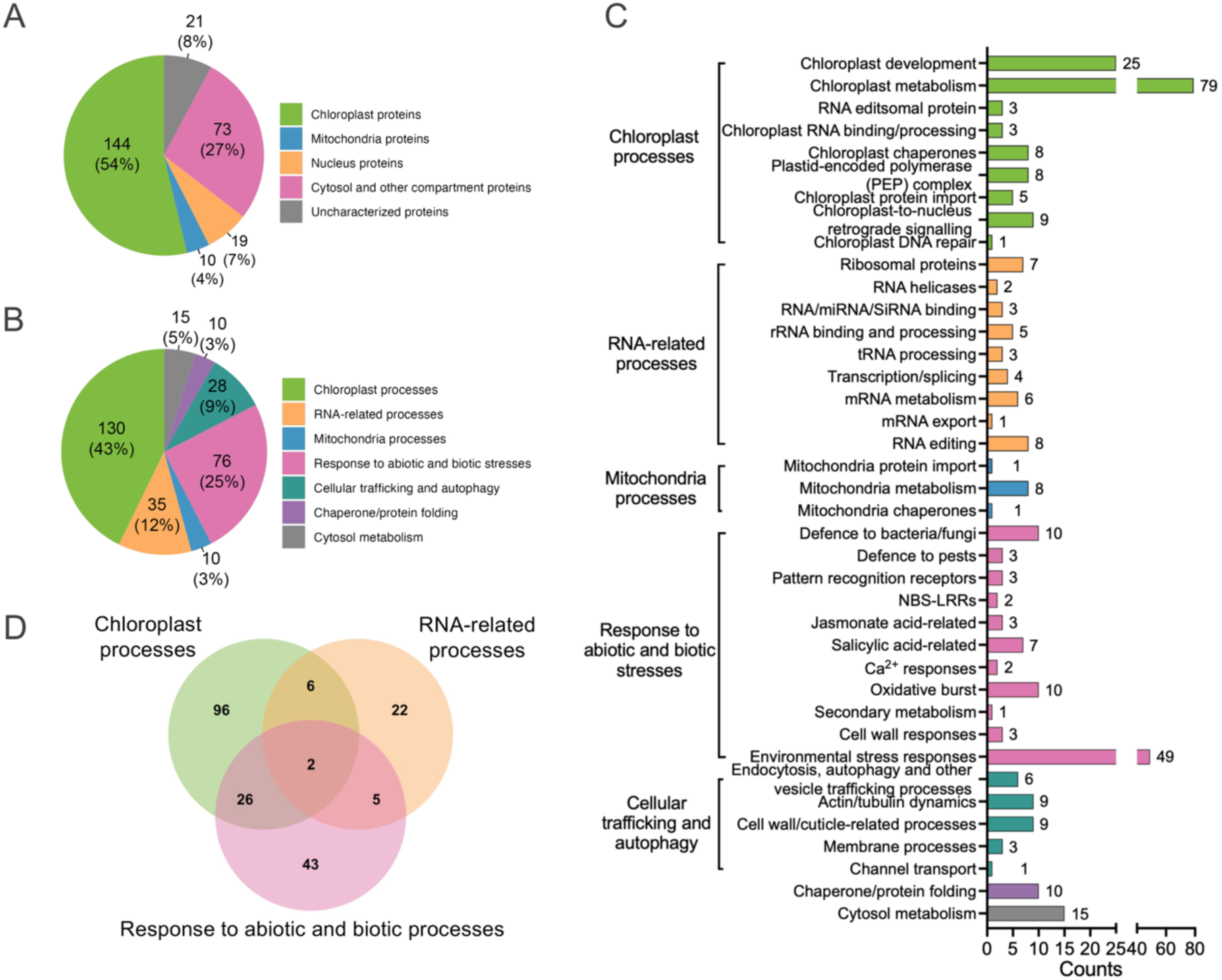
The interactome of *M. persicae* CathB6 in *A. thaliana* identified using TurboID-based proximity labelling. (A) A piechart showing cellular distribution of the biotinylated proteins with the foldchange of CathB6-TurboID/GFP-TurboID > 2 (n = 267). (B) Functional categories of the functionally characterized biotinylated proteins with CathB6-TurboID/GFP-TurboID > 2, shown as piechart. Proteins that were functionally uncharacterized were not included. (C) A barchart showing all functions of characterized biotinylated proteins with CathB6-TurboID/GFP-TurboID > 2. Proteins in different function categories were shown in different color. Protein numbers of each function were labelled on the bar. (D) The overlap of three function categories, chloroplast processes, RNA-related processes and response to abiotic and biotic processes, shown as venn diagram.

### Captions for Supplementary Tables

**Table S1.** List of differentially expressed (DE) genes in plants overexpressing CathB6- GFP vs. GFP empty vector control (GFP), and in *eds1-2* mutant plants vs. wild-type Col-0 controls. Also shown is the overlap with genes detected in comparisons of the *glk1.1 glk2.1* mutant; overexpression of MORF2 (MORF2-OX), and the *gun1-9* mutant relative to respective wild-type control plants from Ahmad et al. (2019) and Zhao et al. (2019). Note that MORF2-OX, *gun1-9* mutants and respective wild-type controls are treated with the herbicide norflurazon.

**Table S2.** Gene Ontology (GO) analysis of over-represented functional categories of up- and down- regulated DE genes for the CathB6-GFP vs. GFP comparison. The table lists the Biological Process GO, group sizes, observed vs expected numbers of DE genes, fold-enrichment and *P*-values.

**Table S3**. GO analysis of over-represented functional categories of up- and down-regulated DE genes for the *eds1-2* vs. wild-type Col-0 comparison. The table lists the Biological Process GO, group sizes, observed vs expected numbers of DE genes, fold-enrichment and *P*-values.

**Table S4**. *A. thaliana* proteins identified as interactors in a Y2H screen of *M. persicae* CathB6 against a cDNA library of *A. thaliana* colonized by *M. persicae*, other insects and a phytoplasma. Plasmids were isolated from yeasts and sequenced. The start and end position relatively to the CDS is indicated for full length (F) and fragments (P).

**Table S5**. Proteins significantly enriched in CathB6-TurboID samples relative to GFP-TurboID controls, identified by proximity labelling-mass spectrometry (PL-MS) in *A. thaliana*. Enrichment was defined as a >2-fold change (CathB6-TurboID/GFP-TurboID) with *P* < 0.05 (two-tailed Student’s t-test). A total of 267 *A. thaliana* proteins met these criteria.

**Table S6**. Gene ontology enrichment analysis of *A. thaliana* proteins identified in CathB6 proximity labeling mass spectrometry.

**Table S7**. Functional classification of *A. thaliana* proteins identified by CathB6 proximity labeling mass spectrometry.

**Table S8**. *A. thaliana* lines used in the study.

**Table S9**. Plasmids used in the study.

## Notes

https://zenodo.org/records/14181687

### Competing Interest Statement

The authors have declared no competing interest.

